# Recognition of Ovarian Tumor-Derived Non-canonical Peptide Induces Memory-Like Features in Natural Killer Cells

**DOI:** 10.1101/2025.09.15.676162

**Authors:** Yizhe Sun, Yaroslav Kaminskiy, Rui M. Branca, Shuhan Li, Okan Gultekin, Kovi Govindajaran, Sahar Salehi, Janne Lehtiö, Dhifaf Sarhan

## Abstract

Immune cell-based immunotherapy has emerged as a promising strategy for both hematologic malignancies and solid tumors. The adaptive properties of T and B cells, namely, antigen specificity and long-term immune memory, form the foundation of approaches such as chimeric antigen receptor (CAR) T-cell therapy and tumor vaccines. In contrast, natural killer (NK) cells, traditionally classified as innate lymphocytes, have been appreciated primarily for their immediate cytotoxicity against tumor cells, but not for long-term memory-like responses. Recent evidence has revealed a subset of NK cells defined as adaptive NK cells (aNK) capable of developing adaptive features, particularly in response to cytomegalovirus (CMV) primarily, and more recently, to ovarian tumor-derived antigens. However, whether tumor-derived peptides can specifically induce NK cell memory, and the corresponding interaction patterns, remains largely unexplored.

In this study, we employed a proteogenomic approach combining RNA sequencing (RNA-seq) with HLA-E immunoprecipitation to identify both canonical and non-canonical peptides presented by primary ovarian tumor cells. Among four tumor-derived neo-antigenic peptides, the 9-mer peptide APAPAPAPL demonstrated the strongest binding affinity to HLA-E and engagement with the NK receptors NKG2C/A. Functional *in vitro* assays confirmed that this peptide could induce memory-like NK cell responses, including antigen-specific recall activity and enhanced tumor cytotoxicity. Furthermore, structural modeling using AutoDock Vina and Rosetta Dock illustrated that peptides with similar binding capacity shared conserved interaction patterns and docking orientations.

Together, this systemic study highlights a novel mechanism for inducing NK cell memory through tumor-derived neoantigens. It also paves the way for the development of NK cell-targeted cancer vaccines, representing a new direction in tumor immunotherapy beyond conventional T cell-centered strategies.

## Introduction

Adaptive immunity is a fundamental component of the human immune system, characterized by long-lasting memory, antigen-specific responses, and enhanced reactivity upon re-exposure. It is established through a relatively well-defined process involving primary antigen exposure and subsequent re-encounters with the same antigens. This process is orchestrated by polymorphic antigen receptors—such as T cell receptors (TCRs) and B cell receptors (BCRs, i.e., membrane-bound immunoglobulins)—as well as peptide-presenting ligands such as major histocompatibility complex (MHC) molecules or intact antigens in the case of BCRs^[1, 2]^. Together, these elements ensure broad recognition of diverse antigens, including tumor-specific antigens (TSAs) and tumor-associated antigens (TAAs), which have already become the key targets to discover^[3, 4]^.

Leveraging this adaptive mechanism, tumor immunotherapy aims to enhance immune responses or engineer immune cells with tailored antigen specificity, based on which two major strategies have emerged: monoclonal antibody-based checkpoint blockade^[5, 6]^ and adoptive cell therapy using immune cells armed with tumor-targeting receptors^[7]^. A complementary approach involves tumor antigen-based vaccines in form of processed peptides or mRNA^[8, 9]^, which seek to elicit both prophylactic and therapeutic immunity by activating and training T and B cells ^[10]^. Encouraging advances regarding tumor vaccines have been achieved in various cancers, including pancreatic cancer, renal cell carcinoma and melanoma^[11-13]^.

Immunotherapy strategies have thus far largely focused on harnessing adaptive immunity. However, chimeric antigen receptors (CARs) have also been engineered into innate immune cells, broadening the therapeutic landscape. CAR-macrophages (CAR-M) exhibit enhanced tumor infiltration, direct phagocytic activity, and intrinsic antigen processing and presentation capabilities, thereby serving as a bridge between innate and adaptive immunity^[14, 15]^.

Promising outcome has emerged in HER2 -overexpressing advanced breast solid tumors^[16, 17]^. Similarly, CAR-engineered natural killer (CAR-NK) cells offer distinct advantages^[18, 19]^, including improved safety profiles, off-the-shelf applicability, and innate cytotoxicity mediated through a balance of activating and inhibitory receptors. Hematological malignancies have emerged as a responsive target for CAR-NK therapy^[20]^. Consequently, NK cells are increasingly recognized for their therapeutic potential in cancer immunotherapy^[21]^.

Despite their therapeutic potential, several limitations and hurdles hinder the application of NK cells in tumor immunotherapy. Unlike adaptive immune cells, NK cells are traditionally classified as part of the innate immune system, owing to their MHC-unrestricted cytotoxicity and dependence on a repertoire of germline-encoded activating and inhibitory receptors^[21, 22]^. As a result, NK cells have long been considered to lack antigen specificity, long-term persistence, and memory capacity, reflecting their relatively limited lifespan *in vivo*^[19]^.

However, accumulating evidence has identified a subset known as adaptive NK (aNK) cells, which exhibit memory-like properties. This phenomenon was first observed in response to murine cytomegalovirus (CMV) infection ^[23-25]^ and has since advanced NK cell biology to overcome its inherent drawbacks for canonical NK cells . and simultaneously challenged the conventional dichotomy between innate and adaptive immunity.

Subsequent studies have expanded the understanding of NK cell memory and its underlying mechanisms. In addition to murine CMV, human cytomegalovirus (HCMV) has also been shown to induce memory-like NK cell responses, and other viral infections such as human immunodeficiency virus (HIV) and simian immunodeficiency virus (SIV) have similarly been implicated in the generation of NK cell memory^[26, 27]^. Furthermore, similar findings have revealed that NK cell memory can exhibit antigen specificity, suggesting that specific processed peptides may contribute to the formation and maintenance of memory-like NK cell responses. Distinct peptides elicit differential recognition by NK cells, ultimately shaping memory formation^[28]^. This antigen-specific recognition has been shown to rely on the activating NK receptor NKG2C, an important marker for aNK cells in human. HLA-E, a non-canonical MHC-I molecule, presents peptides and interacts with NKG2C as a ligand ^[29, 30]^. Although viral antigens can be presented and act as triggers for NK cell memory, whether tumor antigens, especially neoantigens, can elicit a similar response remains unclear, especially considering the potential application of aNK cell in tumor immunotherapy.

Our previous study demonstrated that aNK cells can exhibit tumor-specific memory toward ovarian cancer using tumor cell lysates^[31]^. In this study, we push forward to find the specific tumor peptides derived from the ovarian tumors as the tumor antigens. Commonly, tumor neoantigens are commonly identified through integrative genomic approaches that combine mutation calling with HLA binding prediction. However, direct identification using HLA immunoprecipitation and mass spectrometry offers a more accurate reflection of naturally presented peptides. These methods, together with functional assays, help uncover immunogenic tumor antigens for targeted immunotherapy^[32, 33]^.

Here, we applied a similar workflow focusing on HLA-E, a key ligand for aNK cells. Using primary ovarian tumor cells, we identified both canonical and non-canonical HLA-E-bound peptides. Among four tumor-derived 9-mer candidates, APAPAPAPL showed the strongest HLA-E binding and highest affinity for NKG2C/A receptors. Functional assays *in vitro* confirmed its ability to induce memory-like NK cell responses, including antigen-specific recall and enhanced tumor cytotoxicity. Single-cell transcriptomics revealed distinct gene programs in peptide-educated NK cells, consistent with a memory-like phenotype. Structural modeling further showed that peptides with similar binding strengths shared conserved docking patterns and interaction interfaces.

In summary, this study identifies a tumor-derived peptide that induces memory-like NK cell responses via HLA-E, supporting the therapeutic potential of aNK cells and providing a basis for the development of aNK-targeted tumor vaccines.

## Results

### Proteogenomic analysis and HLA-E binding prediction identified multiple canonical and non-canonical peptides derived from ovarian tumors

Primary ovarian tumor cells (hereafter referred to as **IPLA tumor cells**) were derived from tumor tissues of patients enrolled in a phase III clinical trial, Intra-Peritoneal Local Anesthetics in Ovarian Cancer (IPLA-OVCA) (NCT04065009). The isolated IPLA tumor cells were evaluated both morphologically **(Supplementary Figure 1A)** and phenotypically using the epithelial marker pan-cytokeratin (Pan-CK) and the fibroblast marker S100A4 **(Supplementary Figure 1B and C)**. The markedly higher expression of Pan-CK relative to S100A4 confirmed their epithelial identity and validated them as authentic primary tumor cells.

To comprehensively profile the HLA-E peptide repertoire presented by primary ovarian tumor cells, we performed HLA-E-specific immunoprecipitation followed by liquid chromatography-tandem mass spectrometry (LC-MS) analysis from seven independent patient samples. In parallel, RNA-seq was conducted to support peptide identification and classification for 4 patients. The identified spectra were searched against both canonical and six-frame translated transcriptomic databases using PEAKS Studio and GibbsCluster to refine peptide prioritization **(Figure 1A)**.

**Figure 1.**
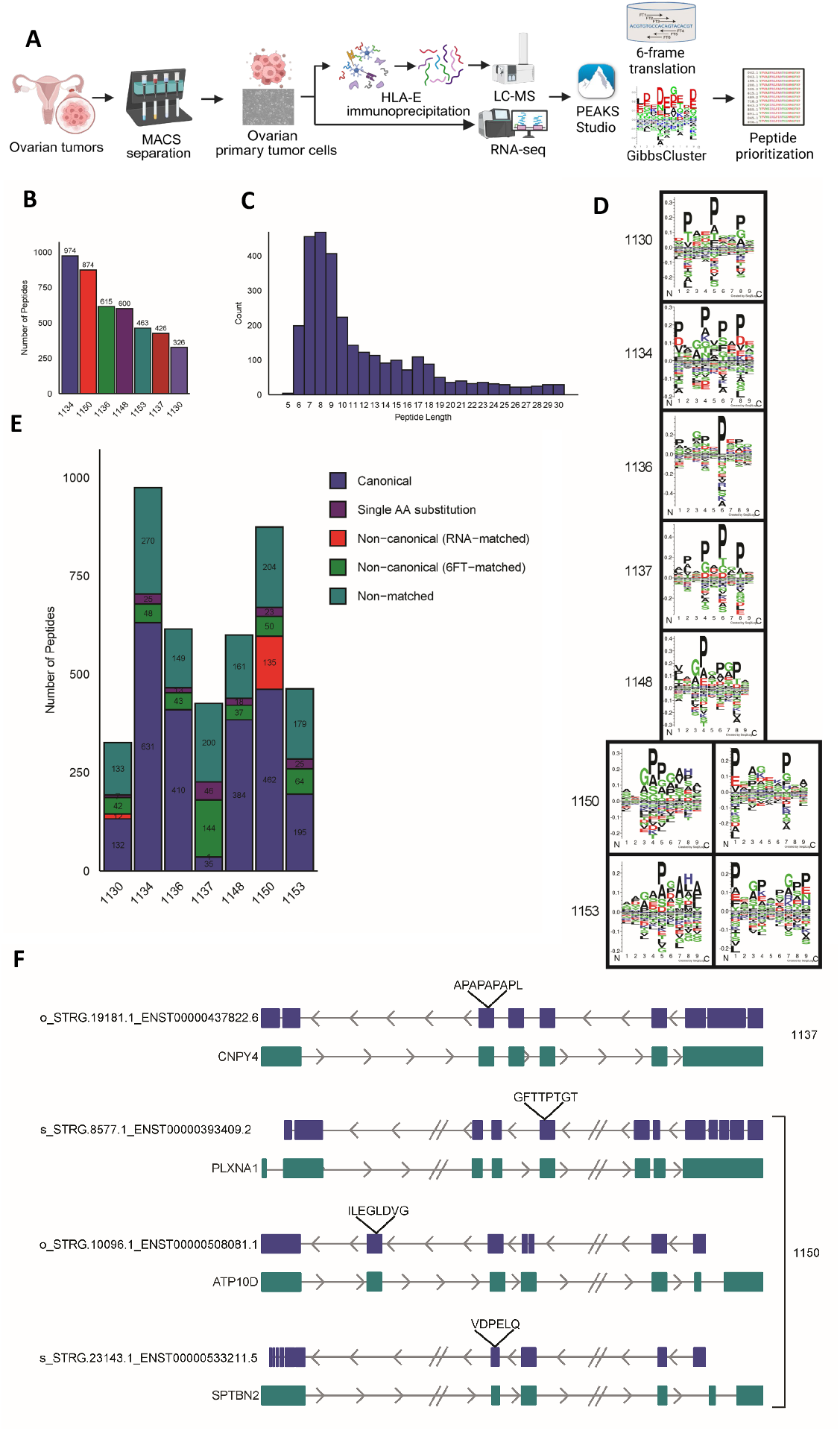
Workflow and characterization of HLA-E–presented peptides in ovarian tumors. **(A)** Experimental workflow: ovarian tumor samples were dissociated, tumor cells isolated by MACS, and HLA-E ligands purified by immunoprecipitation followed by LC-MS/MS analysis and bioinformatic prioritization. **(B)** Number of unique peptides identified from individual patients. **(C)** Length distribution showing enrichment of 8–12mer peptides. **(D)** Sequence motifs from representative patients, highlighting proline enrichment at central positions. **(E)** Categorization of peptides into canonical, single amino acid substitutions, non-canonical (RNA- or six-frame– matched), and non-matched groups across samples. **(F)** Representative non-canonical peptides (APAPAPAPL, GFTTPTGT, ILEGLDVG, VDPELQ) mapped to genomic loci, indicating their derivation from alternative translation or non-canonical transcripts.

A total of 328 to 874 peptides were identified across individual tumor samples **(Figure 1B)**, with a dominant peptide length distribution centered around 9 to 11 amino acids **(Figure 1C)**, consistent with typical HLA-E ligand characteristics. Also, sequence logos of 9-mer peptides across different patients revealed conserved anchor motifs at P2 and P9, but also displayed heterogeneity suggestive of patient-specific preferences **(Figure 1D)**. Peptides were further classified into five categories based on their sequence origin and alignment: canonical, single amino acid substitution, non-canonical (RNA-matched), non-canonical (6-frame translation-matched), and unmatched (middle bottom). Notably, a substantial fraction of peptides (up to ∼40%) fell into non-canonical or unmatched categories, indicating a previously underappreciated diversity in the HLA-E ligandome in tumors **(Figure 1E)**. The final selection of specific peptides was based on stringent criteria: 100% amino acid confidence, an exact match to reference databases, and a peptide length between 6 and 15 amino acids.

This selection encompassed both canonical and non-canonical peptides identified across all seven patients **(Supplementary Table 1)**. To assess the conservation and individuality of the HLA-E-presented peptides, we compared the repertoires across seven ovarian tumor samples. An UpSet plot showed that majority of peptides were patient-specific, while a subset was shared among multiple patients. A corresponding bar plot further confirmed that on average, over 70% of peptides were unique to individual tumors, highlighting both the diversity and partial overlap of the HLA-E ligandome. These findings suggest the presence of tumor-specific HLA-E ligands, which may inform personalized immunotherapy **(Supplementary Figure 2A)**.

Representative examples of non-canonical peptides as novel peptides derived from non-canonical open reading frames (ORFs), including APAPAPAPL, GFTTPTGT, ILEGLDVG, and VDPELQ, were mapped to their transcript structures (bottom panel), further supporting their authenticity and origin from tumor-specific transcripts or alternative translation events. Specifically, APAPAPAPL was derived from an unannotated ORF within the 3’ UTR of *CNPP4*, supported by RNA expression in patient 1137. GFTTPTGT originated from an upstream ORF (uORF) within *PLXNA1* and was detected only in patient 1150, where transcript abundance was also confirmed. ILEGLDVG mapped to an internal out-of-frame region of *ATP10D*, while VDPELQ was derived from a 5’ extended region of *SPTBN2*, both supported by RNA-seq reads and observed exclusively in tumor samples with matching expression **(Figure 1F)**.

To validate the transcriptomic origin of the selected candidate peptides, we examined RNA-seq expression profiles of their corresponding transcripts (*CNPY4, PLXNA1, ATP10D*, and *SPTBN2*) in tumor samples. All four genes were detected with measurable transcript abundance (TPM > 1) across patients, with percentile ranks placing *PLXNA1* and *SPTBN2* in the relatively high range, whereas *CNPY4* and *ATP10D* exhibited slightly lower to intermediate expression. These findings confirm that the identified non-canonical peptides are derived from *bona fide* transcribed genes in ovarian tumors, supporting their presentation as HLA-E ligands **(Supplementary Figure 2B-E)**.

Together, these findings reveal a highly diverse and partially tumor-specific HLA-E ligandome in ovarian cancer, with a substantial portion of peptides derived from alternative or non-canonical sources. This suggests that HLA-E may present a broader repertoire of tumor-derived antigens than previously appreciated, opening new avenues for antigen discovery and immunotherapeutic targeting.

### Interactions between the peptides and both HLA-E and NKG2C/A were identified

Following the prediction of peptides derived from IPLA primary ovarian tumor cells based on their potential to bind HLA-E, we synthesized the 4 non-canonical peptides. To evaluate their binding to HLA-E, we utilized a K562 cell line overexpressing HLA-E*0101 (**K562E**) and performed HLA-E stabilization assays. After testing a range of peptide concentrations **(Supplementary Figure 3A)**, we selected 100 μM and 500 μM for downstream assays. All five candidate peptides, along with the control CMV UL40 peptide (**VMAPRTLIL)**, enhanced surface HLA-E expression, with APAPAPAPL inducing levels comparable to the UL40 peptide **(Figure 2A, 2B)**.

**Figure 2.**
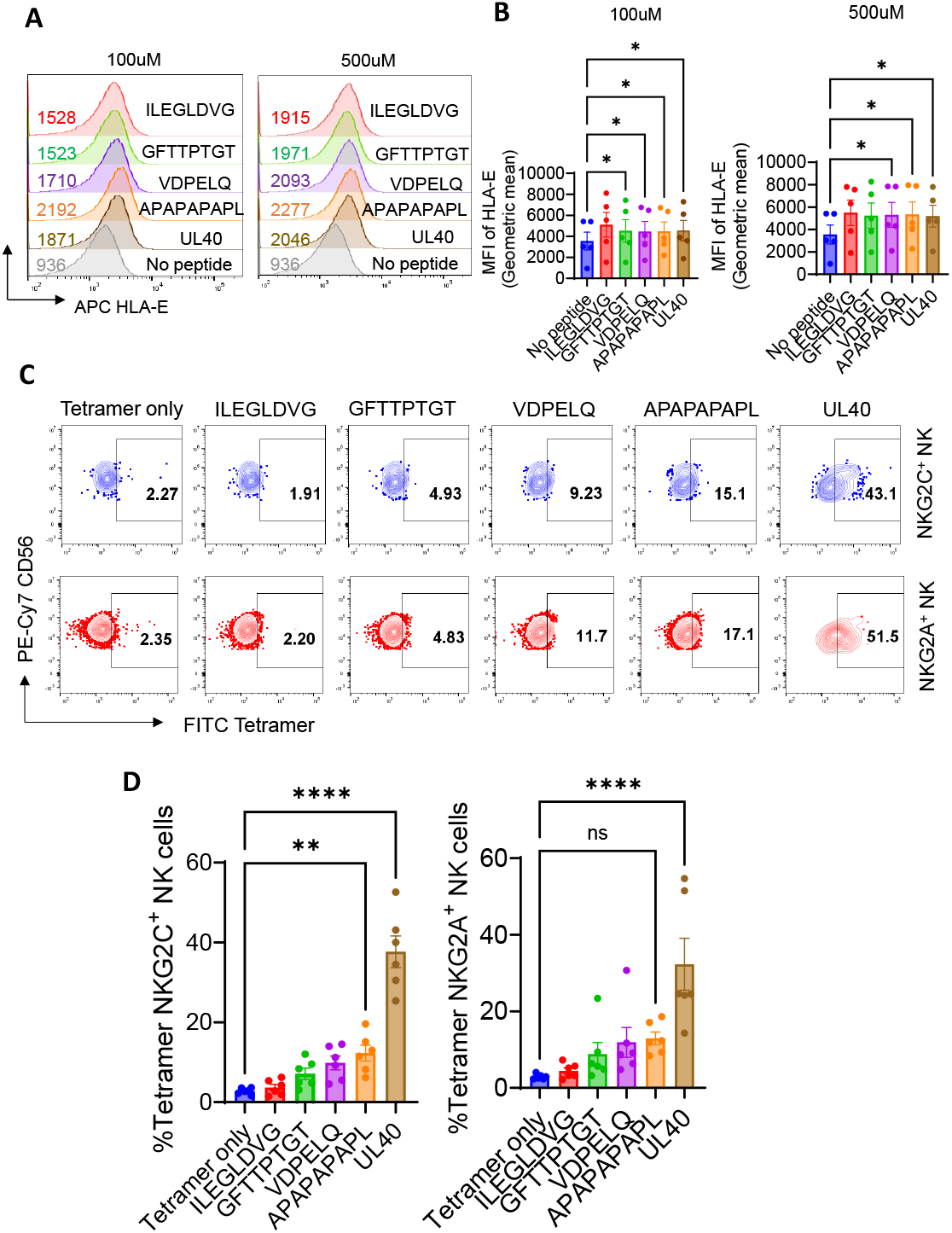
Tumor-derived non-canonical peptides stabilize HLA-E and bind to NK cells. **(A)** Representative flow cytometry histograms showing stabilization of HLA-E on K562-HLA-E*0101 cells loaded with non-canonical peptides (ILEGLDVG, GFTTPTGT, VDPELQ, APAPAPAPL) compared with no peptide controls. **(B)** Quantification of surface HLA-E expression and mean fluorescence intensity across peptides (n=5). Accumulated data are shown as mean ± SEM and statistical analyses using a one-way ANOVA test (*p < 0.05) were conducted using no peptide group as a control. **(C)** Representative tetramer staining of NKG2C^+^ and NKG2A^+^ NK cells, showing binding of HLA-E tetramers loaded with different peptides. **(D)** Summary plots of tetramer-positive NKG2C^+^ and NKG2A^+^ NK cell frequencies across donors, demonstrating selective recognition of tumor-derived non-canonical peptides (n=6). Accumulated data are shown as mean ± SEM and statistical analyses using a one-way ANOVA test (**p < 0.01, ****p<0.0001) were conducted using tetramer only (without peptide loaded) group as a control.

Subsequently, we examined whether the HLA-E-peptide complexes could be recognized by NKG2C^+^ or NKG2A^+^ NK cells. For this purpose, HLA-E tetramers (allele: *0103) loaded with the respective peptides were used to assess their binding to NK cell subsets. The gating strategy for NKG2C^+^ and NKG2A^+^ NK cells was shown **(Supplementary Figure 3B)**. We observed that both NKG2C^+^ and NKG2A^+^ NK cells were capable of binding the HLA-E-APAPAPAPL and HLA-E-VMAPRTLIL complexes, albeit with lower intensity compared to the UL40-loaded complex. Notably, APAPAPAPL appeared to form functional ligands for NKG2C/A (**hereafter referred to NKG2x)** when presented by HLA-E, although the binding of APAPAPAPL to NKG2A^+^ cells did not reach statistical significance despite exhibiting a similar trend **(Figure 2C, 2D)**.

Interestingly, together with the results above, the key variability in NK cell recognition seemed to arise not only from differential peptide-HLA-E binding, but from the differential engagement of NKG2x receptors by the resulting peptide-HLA-E complexes. This suggests that while several peptides can similarly stabilize HLA-E, their capacity to engage NKG2A or NKG2C varies considerably especially for UL40 peptide.

### The peptides serve as the primary immunogenic stimulus for naïve aNK cells

Since we had previously demonstrated that the APAPAPAPL peptide can bind to both HLA-E and the activating NK cell receptor NKG2C, we next sought to investigate its immunogenic potential in the context of dendritic cell (DC)-mediated antigen presentation. As DCs are key initiators of innate and adaptive immune responses, they represent the primary cellular interface for peptide recognition in peripheral tissues. To model this, we pulsed DCs with individual peptides and co-cultured them with NK cells freshly isolated from autologous peripheral blood mononuclear cells (PBMCs), representing a naïve NK cell pool prior to peptide exposure.

Compared to DCs without peptide loading, peptide-pulsed DCs triggered substantial activation of aNK cells after 6h coculture with DCs loaded with the peptides, **(Gated by Supplementary Figure 4A)**. This activation was functionally characterized by a significant upregulation of CD107a (a marker of degranulation), increased secretion of IFNγ and TNFα, and enhanced proliferation as indicated by Ki67 expression. Among all four non-canonical peptides tested, APAPAPAPL consistently induced the strongest primary stimulation across all functional parameters **(Figure 3A, 3B)**, indicating its potent immunogenic capacity.

**Figure 3.**
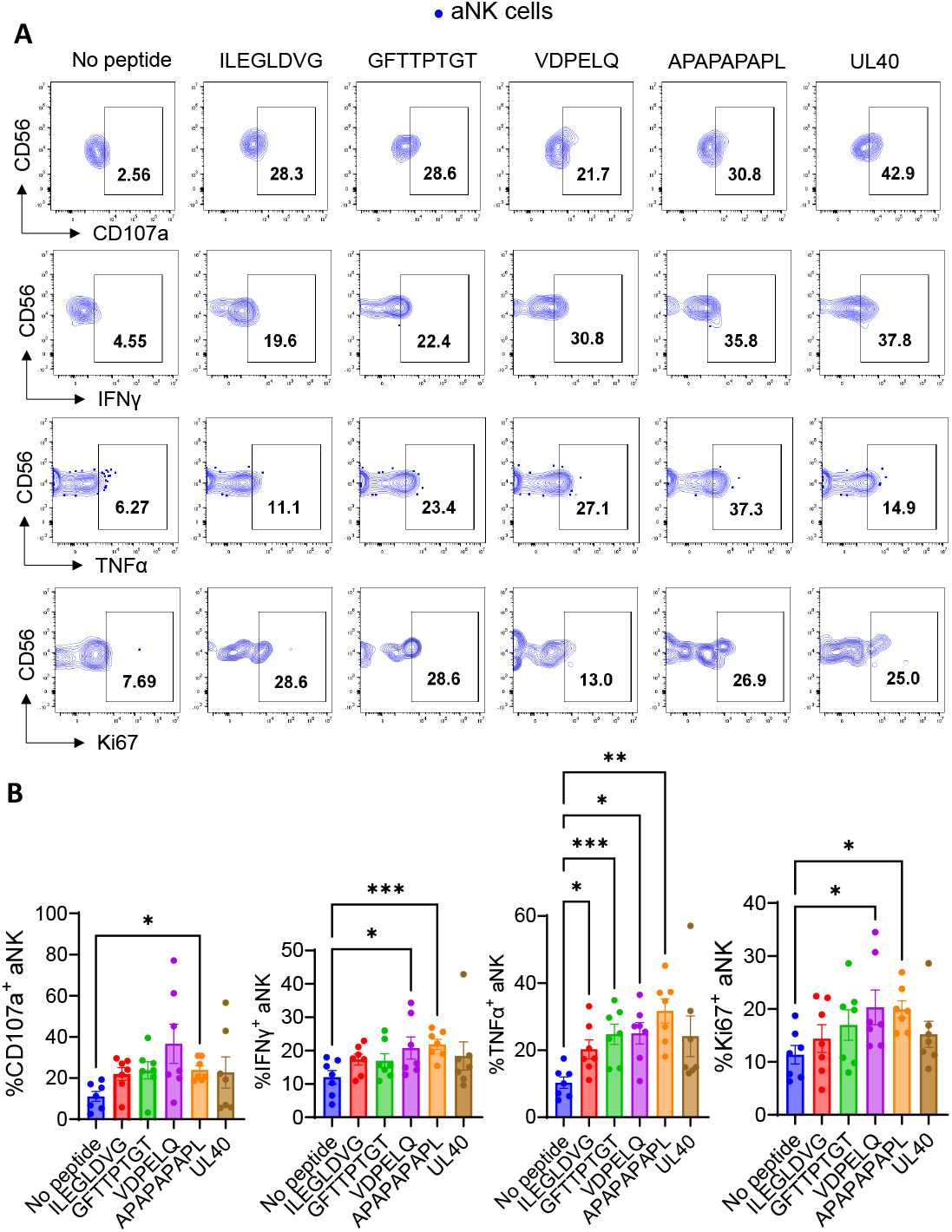
Tumor-derived peptides loaded on DCs trigger primary NK cell responses. **(A)** Representative flow cytometry plots showing aNK cell functional readouts after 6 h co-culture with autologous DCs pulsed with the indicated peptides in terms of degranulation, cytokine production and proliferation (CD107a, IFNγ, TNFα, and Ki67) . **(B)** Summary of aNK cell degranulation, cytokine production and Ki67 expression across donors, demonstrating that tumor-derived non-canonical peptides can elicit primary aNK cell responses (n=7). Accumulated data are shown as mean ± SEM and statistical analyses using a one-way ANOVA test (*p < 0.05, **p<0.01, ***p<0.001) were conducted using no peptide group as a control.

Moreover, when comparing responses between aNK cells and conventional NK (cNK) cells, aNK cells displayed markedly enhanced reactivity to peptide-loaded DCs. This was reflected in higher levels of degranulation, cytokine production, and proliferative activity across the board **(Supplementary Figure 4B)**, suggesting that aNK cells are preferentially responsive to peptide-specific stimulation and may possess an intrinsic capacity for immunological memory-like behavior.

Taken together, these results demonstrate that DC-mediated presentation of non-canonical peptides, particularly APAPAPAPL, can efficiently prime adaptive NK cells, leading to robust functional activation. This highlights the potential role of aNK cells as key effectors in peptide-specific immune responses and suggests that certain tumor-derived or viral peptides may serve as targets for NK cell-based immunotherapeutic strategies. Notably, the pronounced immunogenicity of APAPAPAPL suggests its potential value as a candidate epitope for the development of NK cell-targeted vaccines or memory NK cell activation protocols in cancer or infectious disease settings.

## The peptide induced the enhanced and specific recall responses for aNK cells upon the re-exposure to autologous primary tumor cells

Building on the observation that aNK cells are capable of mounting primary responses to non-canonical peptides, particularly APAPAPAPL. We next investigated whether aNK cells could develop memory-like properties upon repeated peptide exposure. To this end, naïve NK cells were co-cultured with DCs either loaded with peptides or left unpulsed for a period of three weeks, to allow for peptide-driven education.

Following this priming phase, peptide-educated NK cells were restimulated with autologous IPLA primary ovarian tumor cells (the endogenous source of the peptides) or with an unrelated allogeneic pancreatic cancer cell line (CFPAC) serving as a specificity control.

Remarkably, upon restimulation with APAPAPAPL-loaded DCs, aNK cells demonstrated clear recall responses, characterized by elevated degranulation (CD107a expression), increased secretion of effector cytokines (IFNγ and TNFα), and enhanced proliferative capacity (Ki67 expression). These responses were specifically directed against IPLA tumor cells, but not against the irrelevant CFPAC cells, underscoring the antigen-specific nature of the memory-like activation Furthermore, these recall responses were substantially stronger than the primary responses observed in NK cells that were not previously exposed to peptides, indicating that peptide priming conferred an enhanced, memory-like functional capacity. In contrast, cNK cells failed to exhibit such specificity, displaying minimal differences in response regardless of whether the restimulation was autologous or allogeneic **(Figure 4A, 4B)**.

**Figure 4.**
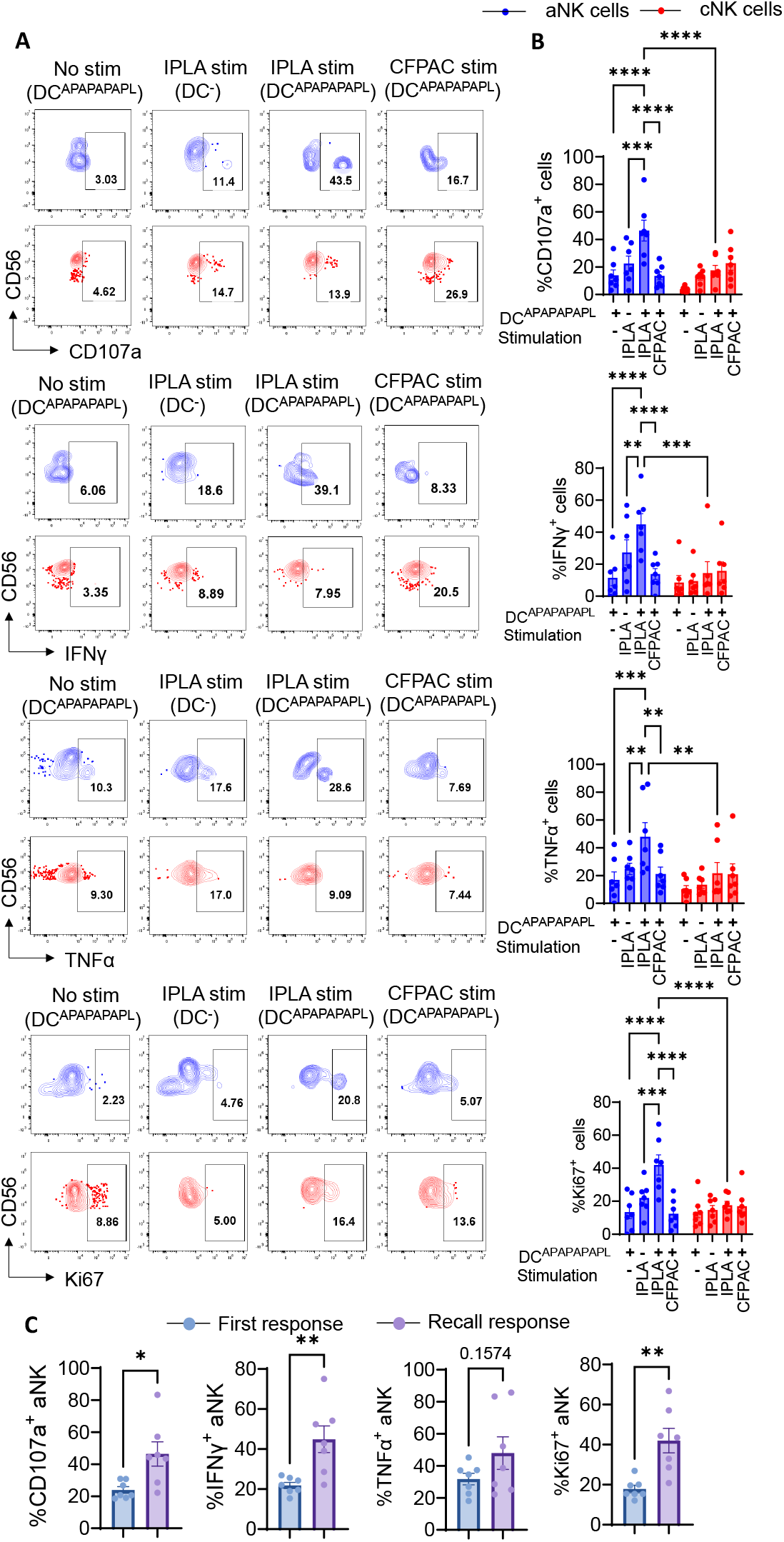
APAPAPAPL-loaded DCs trigger recall NK cell responses. **(A)** Representative flow cytometry plots of aNK and cNK cells upon IPLA tumor restimulation after 3-week co-culture with autologous DCs pulsed with APAPAPAPL, showing degranulation, cytokine production, and proliferation (CD107a, IFN-γ, TNF-α, Ki67). **(B)** Summary of aNK and cNK responses across donors (n = 7). Data are mean ± SEM;statistics by a two-way ANOVA (**p < 0.01, ***p < 0.001, ****p < 0.0001). **(C)** Comparison of primary versus recall responses to APAPAPAPL in aNK cells (n = 7). Data are mean ± SEM; statistics by paired-T test (*p < 0.05,**p < 0.01).

The other non-canonical peptides tested failed to elicit notable recall responses after DC co-culture compared to peptide APAPAPAPL **(Supplementary Figure 5A-5C)**, suggesting suboptimal peptide presentation or insufficient TCR-like recognition by aNK cells during the priming phase. To rigorously define these responses as memory-like, we directly compared the magnitude of APAPAPAPL-induced primary versus recall responses in aNK cells. Recall responses consistently exceeded those of the primary stimulation across all functional parameters **(Figure 4C)**, fulfilling a key criterion for immunological memory.

Collectively, these findings demonstrate that aNK cells, upon DC-mediated peptide education, can develop peptide-specific, memory-like functional responses, with APAPAPAPL emerging as a potent immunogenic epitope capable of inducing durable aNK cell memory-like features.

### Blocking assays identified the NKG2C as the central receptor and HLA-E as the crucial ligand for the recognition of the peptide

To further elucidate the molecular mechanism underlying APAPAPAPL-induced primary and recall responses, we performed a series of blocking assays using neutralizing antibodies against key surface molecules, including NKG2C, NKG2A, HLA-E, MHC-I, and MHC-II.

We first examined the first responses of NK cells upon APAPAPAPL peptide exposure. Neutralization of NKG2C markedly diminished both degranulation (CD107a expression) and IFNγ production in aNK cells and also exhibited a downward trend in TNFα production.

Blocking HLA-E also led to a noticeable reduction trend in degranulation and IFNγ production, suggesting that the APAPAPAPL-mediated primary activation of aNK cells is, at least in part, dependent on NKG2C-HLA-E interactions. In contrast, cNK cells showed minimal or no changes in first responses upon blockade of either NKG2C or HLA-E, indicating a lack of reliance on this interaction axis for initial activation **(Supplementary Figure 6A-6B)**.

To assess the receptor dependency of the recall responses, we next evaluated the effects of blocking during and after the 3-week peptide-based priming. In a restimulation-phase blockade setting, NK cells were treated with neutralizing antibodies during the 6-hour restimulation phase with autologous IPLA primary tumor cells. Under this condition, aNK cells exhibited significantly impaired recall responses when NKG2C or HLA-E were blocked, as evidenced by reduced CD107a mobilization and IFNγ production. These effects were not observed in cNK cells, reinforcing the selective reliance of aNK cells on the NKG2C-HLA-E axis for recall responses **(Figure 5A-5D)**.

**Figure 5.**
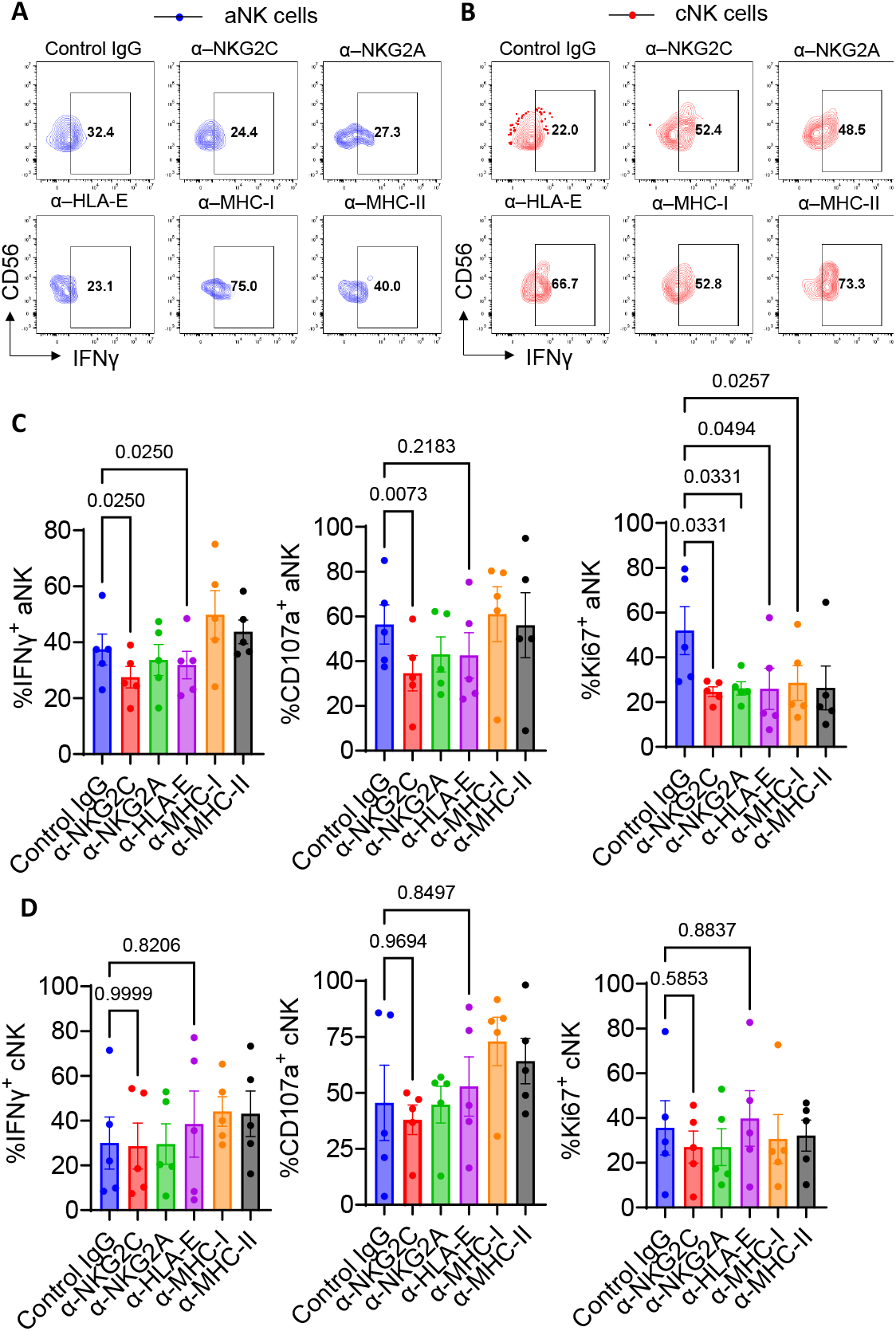
NKG2C and HLA-E are critical for recall responses of aNK cells to APAPAPAPL. **(A–B)** Representative contour plots showing IFN-γ^+^ frequencies in adaptive NK (aNK, A) and conventional NK (cNK, B) cells following re-exposure to IPLA tumor cells, with blocking of the indicated receptors and ligands during the restimulation phase. (C) Statistical summaries of IFN-γ^+^, CD107a^+^, and Ki67^+^ aNK cells after restimulation with blocking antibodies compared to isotype IgG control (n = 5). Data are presented as mean ± SEM and analyzed by one-way ANOVA; p values are indicated in the plots, with all groups compared against the isotype control (**D**) Statistical summaries of IFN-γ^+^, CD107a^+^, and Ki67^+^ cNK cells under the same conditions (n = 5). Data are presented as mean ± SEM and analyzed by one-way ANOVA; p values are indicated in the plots, with all groups compared against the isotype control.

Additionally, in a long-term blockade experiment during the priming phase, neutralizing antibodies were included throughout the entire 3-week coculture with peptide-loaded DCs, thereby disrupting receptor-ligand interactions during both the priming and recall phases. Consistently, long-term blockade of NKG2C profoundly impaired the development of recall responses in aNK cells in terms of IFNγ production and degranulation, blocking of HLA-E also exhibited the similar trend, while cNK cells remained less unaffected, further underscoring the functional specificity of this pathway in memory-like NK cell responses **(Supplementary Figure 6C-6D)**. It demonstrates that the NKG2C-HLA-E interaction is not only essential for triggering the recall response in aNK cells, but also plays a critical role during the initial priming phase necessary for establishing memory-like features.

### *In silico* simulation for the peptide interaction mode with HLA-E and NKG2x

To further investigate the molecular basis of peptide recognition, we conducted a comprehensive series of *in silico* simulations using all peptides identified from our proteogenomic screening including the four non-canonical peptides **(Supplementary Table 1)**. A hierarchical docking workflow was established, integrating AutoDock Vina^[34]^ for initial coarse docking, Rosetta FlexPepDock^[35]^for high-resolution refinement, and RosettaDock ^[36, 37]^ for modeling receptor engagement. Specifically, each peptide was first docked to HLA-E*0103 **(PDB: 1KPR** ^[38]^) using AutoDock Vina to generate preliminary peptide-HLA-E binding poses. The top-ranked model was selected for the four non-canonical peptides with the lowest value of kcal/mol indicating the best affinity **(Supplementary Figure 7A)**. It demonstrated primary APAPAPAPL has the best binding affinity with HLA-E*0103 compared to the other three peptides.

The selected models from AutoDock Vina were then refined using Rosetta FlexPepDock, yielding up to 200 conformers per peptide. These models were evaluated based on total Rosetta energy and reweighted interface scores, and the top five conformations were selected for the overall comparison. Among the four non-canonical peptides analyzed, **APAPAPAPL** consistently demonstrated the most favorable binding characteristics—including the lowest total Rosetta energy, reweighted score, and interface energy—indicating strong and stable interaction with HLA-E **(Figure 6A)**. Principal component analysis (PCA) and K-means clustering on FlexPepDock parameters further revealed that peptides containing conserved structural motifs—such as proline at positions 4 and 6, or a ‘PA’ motif at positions 6-7— tended to cluster with APAPAPAPL. Notably, VMGNNPADL exhibited the closest structural and energetic similarity to APAPAPAPL **(Figure 6B, 6C, Supplementary Figure 7B)**.

**Figure 6.**
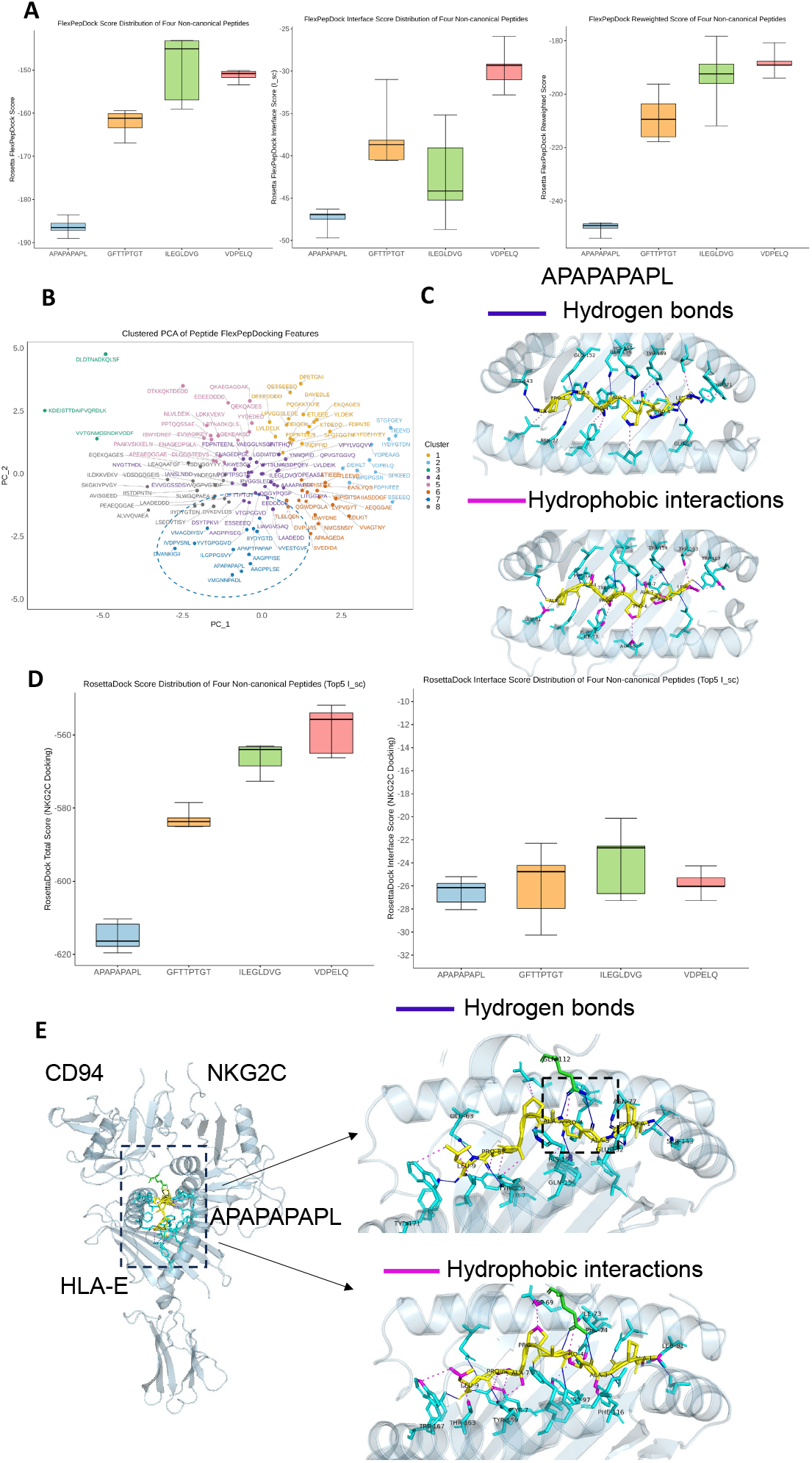
*In silico* simulation for APAPAPAPL interaction with HLA-E and CD94-NKG2C. **(A)** FlexPepDock refinements of APAPAPAPL, GFTTPTGT, ILEGLDVG, and VDPELQ bound to the HLA-E*01:03 complex. Box plots (boxes = interquartile range, line = median, whiskers = range) summarize Rosetta total score, interface ΔG (I_sc), and reweighted score for the top 5 models of each peptide–HLA-E complex. **(B)** PCA mapping of all best models of peptides identified in Figure 1 complexed with HLA-E, showing their clustering based on multiple interaction simulation parameters. **(C)** Visualization of interaction sites and contact types between APAPAPAPL and HLA-E. **(D)** RosettaDock of APAPAPAPL, GFTTPTGT, ILEGLDVG, and VDPELQ-HLA-E complex with CD94-NKG2C complex. Box plots (boxes = interquartile range, line = median, whiskers = range) summarize Rosetta total score and interface ΔG (I_sc) for top 5 I_sc models of each complex. **(E)** Visualization of interaction sites and contact types between APAPAPAPL-HLA-E with CD94-NKG2C complex.

We next modeled the ternary complex formed by HLA-E-peptide with CD94-NKG2C or CD94-NKG2A using RosettaDock, incorporating spatial restraints derived from the reference crystal structure (PDB: 3CDG^[39]^). Among the four non-canonical peptides tested, **APAPAPAPL** consistently yielded the top-scoring models for both NKG2C and NKG2A complexes, exhibiting the lowest interface energies **(Figure 6D, Supplementary Figure 7C)**. In the NKG2C complex, CD94 residue Q112 established multiple hydrogen bonds with alanine residues at positions 3 and 5 of APAPAPAPL, whereas in the NKG2A complex, a single hydrogen bond was observed between Q112 and alanine at position 5 **(Figure 6E)**.

In contrast, peptides such as **ILEGLDVG** and **GFTTPTGT** showed minimal or no interaction with either NKG2C or NKG2A, while **VDPELQ** formed notable salt bridges and hydrophobic interactions with both receptors **(Supplementary Figures 7D-7F)**. Interestingly, **VMGNNPADL** consistently clustered with APAPAPAPL in PCA-based analyses of both NKG2C and NKG2A complexes **(Supplementary Figures 8A-8B)**, suggesting that shared sequence motifs—particularly a “PA” motif and terminal leucine—may play a key role in mediating binding to HLA-E and subsequent engagement with NK cell receptors.

Moreover, comparative analysis of docking metrics revealed that while the interface geometries of NKG2C and NKG2A complexes were largely similar, the total binding energy of the NKG2C-HLA-E-peptide complex was significantly lower (i.e., more favorable), indicating a higher overall stability for APAPAPAPL **(Supplementary Figure 8C)**. These findings support a model in which NKG2A inherently exhibits weaker binding stability toward tumor-derived peptide-HLA-E complexes compared to NKG2C.

Comparative analysis revealed that NKG2C-CD94 generally formed more stable and favorable interactions with the peptide-HLA-E complex than NKG2A-CD94, based on both total energy and interface metrics. These findings highlight APAPAPAPL as a structurally optimal non-canonical peptide for HLA-E presentation and NKG2C recognition, and suggest that specific sequence motifs, particularly proline- and alanine-rich segments, play a critical role in tuning NK cell receptor binding.

## Methods and Materials

### Study approval

Tumor samples from patients with high-grade serous ovarian cancer (HGSOC) were collected between 2020 and 2024 as part of a phase III clinical trial [Intra-Peritoneal Local Anesthetics in Ovarian Cancer (IPLA-OVCA) trial; https://clinicaltrials.gov/ct2/show/NCT04065009]. Eligible patients were enrolled based on a confirmed diagnosis of stage III or IV epithelial ovarian cancer and scheduled for upfront cytoreductive surgery. All participants provided written informed consent prior to inclusion, in accordance with the Declaration of Helsinki. The study protocol was approved by the local Ethics Committee and Institutional Review Board (approval numbers 2019-05149 and 2015/1862-32). Animal experiments conducted in this study were approved by the local Ethics Committee and Institutional Review Board (Dnr 10970-2021), in accordance with institutional and national regulations governing the care and use of laboratory animals. All procedures were performed with strict attention to minimizing animal discomfort and ensuring humane treatment throughout the study.

### Cell lines and primary tumor cells establishment

CFPAC cells (a pancreatic cancer cell line acquired in 2022 from the ATCC) were maintained in standard culture conditions. TYK-nu cells were grown in Minimum Essential Medium α (MEMα; Thermo Fisher Scientific, Cat. #22571020), while CFPAC cells were cultured in RPMI 1640 medium (Thermo Fisher Scientific, Cat. #22400089). Both media were supplemented with 10% heat-inactivated fetal bovine serum (Thermo Fisher Scientific, Cat. #SV10160.03HI) and 1% penicillin-streptomycin (Nordic Biolabs, Cat. #PS-B). Cells were incubated at 37°C in a humidified atmosphere containing 5% CO_2_.

IPLA primary ovarian tumor cells were established from fresh biopsy samples obtained from IPLA trial participants. Tissues were enzymatically digested in harvest medium consisting of RPMI supplemented with 150 μg/mL Liberase (Roche, Cat. #5401127001) and 100 μg/mL RNase I (Roche, Cat. #4716728001) for 30 minutes at 37°C. Following red blood cell lysis (Transfusion Medicine, Karolinska University Hospital), the cell suspension was passed through a 70 μm strainer and then filtered using a 0.8 μm Nalgene Rapid-Flow Sterile Filter Unit (Thermo Fisher Scientific, Cat. #450-0080) to remove residual debris and aggregates.The filtered tumor cells were seeded into T75 flasks and cultured in RPMI medium supplemented with 10% patient-derived ascites. Ascites samples were collected in parallel with tumor tissue and tested for their capacity to support tumor cell outgrowth. The most supportive ascites sample, determined by visual inspection of confluency and viability over time, was selected for continued use. Ascites was heat-inactivated at 56°C for 1 hour prior to use and filtered for sterility. During the monitoring of primary tumor cells, 10% FBS was used for the primary tumor cells culture.

Adherent cells were expanded for 1-2 weeks until reaching ∼90% confluency, at which point tumor cells were selectively enriched using the Tumor Cell Isolation Kit (Miltenyi Biotec, Cat. #130-108-339) following the manufacturer’s protocol. Purity of isolated tumor cells was confirmed via immunofluorescence staining for pan-cytokeratin (PanCK; Abcam, Cat. #ab80826), and the absence of fibroblastic contamination was validated by negative staining for S100A4 using Alexa Fluor 488-conjugated secondary antibodies (Lifespan Biosciences, Cat. #LS-C755562). To minimize mycoplasma contamination, ciprofloxacin hydrochloride (1 μg/mL; GenHunter, Cat. #Q901) was added to the culture medium throughout the process. At each passage, cells were cryopreserved in freezing medium consisting of 90% FBS and 10% DMSO (Sigma-Aldrich, Cat. #276855-100ML).

### HLA-E immunoprecipitation

For immunopeptidomic analysis, an aliquot of purified IPLA primary tumor cells (0.5-1 x 10^7^ cells) was rapidly thawed, washed with PBS and resuspended in 90 µL of PBS. An equal volume (90 µL) of lysis buffer (1% CHAPS, 1x cOmplete™ EDTA-free Protease Inhibitor Cocktail, Roche, P/N 11697498001) was added to the cell suspension, yielding a final concentration of 0.5% CHAPS. Cell lysis was performed for 1 hour at 4°C with gentle end-over-end rotation. The lysate was clarified by sequential centrifugation: first at 2,000g for 10 minutes at 4°C, followed by a high-speed centrifugation of the supernatant at 21,000g for 20 minutes at 4°C. The final supernatant was carefully collected and incubated with anti-human HLA-E antibody (eBioscience™, 3D12HLA-E) immobilized on MagReSyn® Protein A MAX magnetic beads (ReSyn Biosciences, MR-PAM002) overnight at 4°C with rotation as per the manufacturer’s protocol. Following incubation, the beads were washed and HLA-E complexes were eluted with 500 µL of 1% trifluoroacetic acid (TFA). The eluted peptides were separated from the HLA-E complex and antibody using a C18 solid-phase extraction cartridge (Waters™ Sep-Pak® tC18, P/N WAT023501). Peptides were eluted from the C18 cartridge using 40% acetonitrile (ACN) / 0.1% TFA, dried in a vacuum centrifuge, and stored at -80°C until mass spectrometric analysis.

### LC-MS analysis of HLA-E peptides

To each sample dry in a Lo-Bind Eppendorf tube, 20 μL of LCMS solvent A (0.1% formic acid) was added, and after diss the 20 μL were transferred to an Evotip pure (Evosep, Odense, Denmark) and prepared according to the manufacturer’s instructions. The Evosep One LC system (Evosep, Odense, Denmark) then proceeded to inject sample onto the analytical column (Aurora Elite G3, C18, 15 cm long, 75 μm ID, 1.7 μm bead size, Ionopticks, Melbourne, Australia) constantly kept at 50°C by a heating oven (Column Toaster, Bruker, Bremen, Germany) using the Whisper 40 samples per day 31min gradient. The Tims TOF SCP mass spectrometer (Bruker) operated with the CaptiveSpray source, capillary voltage 1500 V, dry gas flow of 3 L/min, dry gas temperature at 180°C. Mass range was 100-1700 m/z, and mobility range was 0.6-1.70 V.s/cm2. MS/MS was used with three PASEF (parallel accumulation - serial fragmentation) scans (with 300 ms ramp time) per cycle with a target intensity of 20 000 and intensity threshold of 500, considering charge states 0-5. Active exclusion was used with release after 0.4 min, reconsidering precursor if the current intensity is greater than fourfold the previous intensity, and a mass width of 0.015 m/z and a 1/k0 width of 0.015 V.s/cm2. Isolation width was defined as 2.00 m/z for mass 700 m/z and 3.00 m/z for mass 800 m/z. Advanced collision energy settings consisted of nine points of collision energy dependency on ion mobility: 0.70 V.s/cm2 - 27.50 eV; 0.85 V.s/cm2 - 32.00 eV; 0.90 V.s/cm2 - 36.00 eV; 0.95 V.s/cm2 - 38.00 eV; 1.25 V.s/cm2 - 44.00 eV; 1.30 V.s/cm2 - 46.00 eV; 1.35 V.s/cm2 - 47.00 eV; 1.50 V.s/cm2 - 47.00 eV; 1.65 V.s/cm2 - 50.00 eV. Precursor ions were selected using 1 MS repetition and a cycle overlap of 1 with the default intensities/repetitions schedule. Selection polygon shape was optimized to increase HLA-E peptides identification number.

### *De Novo* assembly of transcripts from RNA-seq

To support peptide identification and enable the discovery of non-canonical translation events, RNA-seq was performed on matched IPLA primary ovarian tumor cell samples used for HLA-E immunopeptidomics. Sequencing libraries were prepared from 100 ng total RNA, using the Illumina Stranded Total RNA library preparation kit with Ribo-Zero Plus treatment (Cat# 20040525/20040529, Illumina Inc.). Unique dual indexes (Cat# 20040553/20040554, Illumina Inc) were used. The library preparation was performed according to the manufacturers’ protocol (Cat# 1000000124514). Samples were run on NovaSeq X Plus system, with Paired-end 150bp read length, 10B flowcell and XLEAP-SBS sequencing chemistry.

The nf-core/rna-seq 3.8.1^[40]^ workflow was used to pre-process and align the raw reads to GRCh38.p6. Stringtie was used to assemble transcripts from each replicate in reference-guided assembly mode to allow for detection of novel transcripts. Expressed transcript (TPM>1) per sample were obtained by merging the assembled transcripts from all replicates of each sample to obtain a GTF per sample. The transcripts were labeled using gffcompare based on their similarity to the standard transcript database.

### Database search

All MS/MS spectra were searched by PEAKS Studio 11.0 (build 20230414, BSI)^[41]^ using a Deep Novo search (database-free *de novo* search) first, then followed by a Peptide Search (database search with target-decoy strategy). The search database was either a sample-specific FASTA generated by three-frame translation (3FT) of transcripts assembled from bulk RNA-seq of IPLA primary tumor cells, concatenated with the UniProt human reference proteome (and common contaminants), or the UniProt human reference proteome alone. Additionally DDA contamination database was included^[42]^. A precursor mass tolerance of 20 ppm and a fragment ion mass tolerance of 0.05 Da were used. Enzyme was none, digest mode unspecific, and oxidation of methionine was used as variable modification, with max two oxidations per peptide. An FDR cutoff of 5% was employed at the peptide level. All peptides identified by PEAKS and reported in “peptideDb.peptides” is fed into GibbsCluster - 2.0^[43]^.

### Peptide categorization

Peptides that were found by DeepNovo (without exact match to database) were categorized based on their probable origin. These peptides were restricted by selecting peptides with length between 8 and 15 AA for HLA E. Peptides matching to HLA sequences from peptideAtlas and from IEDB were done using blastp-short with a e-value of 10.0^[44]^. The peptides were searched against the human canonical proteome using blastp-short with an e-value of 10.0 to find single amino acid substitutions and matches to the canonical proteome. Peptides with single amino acid mismatch to the canonical proteome was then searched against dbSAP using blastp-short and a e-value of 10.0^[45]^. Peptide with more than 1 mismatch to the canonical proteome was searched against the human six frame translated genome. The six-frame translation was performed on human reference genome build GRCh38.p14 using SeqKit translate^[46]^. Peptide were searched for exact matches to the six-frame translated (6FT) genome using SeqKit grep. Potential *cis*-proteosome catalysed splicing variants were also detected and classified.

### Peptides synthesis and autologous DC-NK cell cocultures

Synthetic peptides were custom-ordered from MedChemExpress (MCE, Monmouth Junction, NJ, USA) at >95% purity. And all the peptides were dissolved with sterile PBS at 10Mm_。_

Monocytes were isolated from PBMCs of healthy donors using CD14 magnetic microbeads (Miltenyi Biotec, Cat. #130-050-201) according to the manufacturer’s instructions. Isolated monocytes were seeded at a density of 1 × 106 cells/mL in CellGro serum-free DC medium (CellGenix, Cat. #20801-0500) supplemented with 10% fetal bovine serum (FBS) and 1% penicillin-streptomycin (P/S). Differentiation into immature dendritic cells (DCs) was induced by culturing cells in the presence of recombinant human GM-CSF (100 ng/mL; PeproTech, Cat. #300-03) and IL-4 (20 ng/mL; PeproTech, Cat. #200-04) for 48 hours.

Immature DCs were gently detached using a nonenzymatic cell dissociation buffer (Thermo Fisher Scientific, Cat. #13151-014); any remaining adherent cells were recovered by gentle scraping. Maturation was induced by culturing cells in the presence of IL-4, GM-CSF, poly I:C (20 μg/mL; Sigma-Aldrich, Cat. #P1530), resiquimod (R848; 2.5 μg/mL; InvivoGen, Cat. #tlrl-r848), IFNγ (1,000 IU/mL; PeproTech, Cat. #300-02), and lipopolysaccharide (LPS; 10 ng/mL; Sigma-Aldrich, Cat. #437627) for an additional 24 hours. Maturation was carried out in CellGro medium supplemented with 2% heat-inactivated human AB serum (Stockholm Blood Center) and 1% P/S.

Simultaneously, DCs were pulsed with the tumor peptides at a concentration of 10uM. For coculture experiments, mature antigen-loaded DCs were incubated with NK cells in round-bottom 96-well plates using RPMI complete medium supplemented with 10 ng/mL recombinant human IL-15 (Miltenyi Biotec, Cat. #130-095-760). Cocultures were maintained for up to 4 weeks, with IL-15-containing medium refreshed every 7 days.

At weeks 3, NK cells were restimulated using autologous IPLA cells or allogeneic tumor cells (CFPAC) at an effector-to-target (E:T) ratio of 1:1. After restimulation, flow cytometry staining was performed. In long-term blocking experiments, functional blocking antibodies (5 μg/mL) were added either at the initiation of coculture (days 0, 7, and 14) or during the restimulation phase (day 21, for 6 hours). Antibodies included anti-NKG2C (R&D Systems, Cat. #MAB138, clone 134591), anti-NKG2A (BioLegend, Cat. #375102, clone S19004C), anti-HLA-E (BioLegend, Cat. #342602, clone 3D12), anti-MHC class I (Sigma-Aldrich, Cat. #SAB4700633, clone MEM-123), anti-MHC class II (BioLegend, Cat. #365602, clone QA19A44), and isotype-matched control IgG (R&D Systems, Cat. #MAB002, clone 133303).

### Analysis of cytokine production and degranulation by flow cytometry

Proinflammatory cytokine production (IFNγ and TNFα) and degranulation (CD107a expression) were assessed in DC-NK coculture systems using flow cytometry. After DC-NK coculture, NK cells were restimulated for 6 hours with autologous viable tumor cells (IPLA or TYK-nu) or an irrelevant control tumor cell line (CFPAC) at a 1:1 effector-to-target (E:T) ratio. In short-termed blocking assays, functional blocking antibodies abovementioned (5 μg/mL) were added during the 6-hour restimulation Restimulation was performed in RPMI 1640 medium containing anti-CD107a antibody and protein transport inhibitors GolgiPlug (BD Biosciences, Cat. #51-2301KZ; 1:1,000) and GolgiStop (BD Biosciences, Cat. #51-2092KZ; 1:1,000). After stimulation, cells were stained for surface markers, then fixed and permeabilized using the eBioscience Foxp3/Transcription Factor Fixation/Permeabilization kit according to the manufacturer’s instructions. Intracellular staining was performed for markers including FcεRIγ, Ki-67, IFNγ, and TNFα, using antibodies listed in **Supplementary Table 4**. Data acquisition was carried out using a CytoFLEX cytometer (Beckman Coulter), and analysis was performed with FlowJo software (version 10.5).

### Tetramer staining of primary human NK cells

PBMCs were isolated from healthy blood donor buffy coats (Stockholm Blood Center) via density gradient centrifugation using Ficoll-Paque PLUS (GE Healthcare). Bulk NK cells were enriched from PBMCs using the NK Cell Isolation Kit (Miltenyi Biotec, Cat. #130-092-657) according to the manufacturer’s protocol.

For tetramer staining, a pre-assembled FITC-conjugated HLA-E*01:03&B2M tetramer (KACTUS, Cat #MHC-HM42RTR) loaded with peptides was used. The tetramer-peptide complex was freshly prepared at a molar ratio of 20:1 (peptide:tetramer), and 0.1 μg of tetramer was used per 1×10^5^ cells. Isolated NK cells were incubated with the tetramer for 30 minutes at room temperature in the dark.

Following tetramer binding, cells were washed and stained with surface antibodies including anti-CD3 (BV510), anti-CD56(PE-Cy7), anti-NKG2C(PE), and anti-NKG2A (APC) for an additional 20 minutes at 4°C. After washing, cells were acquired using a CytoFLEX flow cytometer and analyzed with FlowJo software (version 10.5).

### Protein and peptide interaction simulations and docking

Peptide docking and receptor complex modeling were performed using a multi-step computational pipeline. First, the HLA-E*0103 heavy chain structure was extracted from the PDB entry 1KPR and prepared by removing existing ligands (peptides) and adding the hydrogens. Candidate peptides were docked into the peptide-binding groove using AutoDock Vina (v1.2), with docking parameters optimized for short peptide-MHC interactions.

The top-ranked binding poses for each peptide were further refined using Rosetta FlexPepDock in high-resolution mode. Each peptide-HLA-E complex was subjected to 200 refinement trajectories, and resulting models were evaluated based on total energy score and interface energy.

To model the interaction between the HLA-E-peptide complex and its receptor CD94-NKG2x, RosettaDock (v3.12) was used in local docking mode. The pre-docked HLA-E-peptide structure was treated as a rigid body (chains H and P), while CD94 and NKG2C/A (chains C and N) were docked against it. A total of 500 docking trajectories were generated per peptide using the -dock_pert 3 8 setting to introduce translational and rotational perturbations around the starting orientation.

To maintain the correct binding orientation and avoid flipping of the complex during sampling, a distance constraint was applied between residue 163 (Cα atom) of HLA-E and residue 211 (Cα atom) of NKG2C/A. This was implemented using a bounded constraint: AtomPair CA 163H CA 211N BOUNDED 3.0 30.0 0.5 no This constraint enforces the interatomic distance to remain between 3 Å and 30 Å, with deviations penalized during energy minimization. The constraint file was provided via the - constraints:cst_file option, and its weight was set to 0.5 using -constraints:cst_fa_weight.

Scoring was performed using the ref2015 energy function. All docking models were saved in the specified output directory, and scorefiles were collected for further ranking and analysis.

### PCA and K-means clustering based on docking energy features

To explore similarities among different peptide-HLA-E-NKG2x docking models from RosettaDock and different peptide-HLA-E docking from FlexPepDock, we selected the top-ranked conformation (i.e., lowest interface energy) of each peptide and performed PCA followed by K-means clustering.

For RosettaDock outputs, 13 energy terms were extracted for PCA, including overall energy (total_score), interface score (I_sc), van der Waals terms (fa_atr, fa_rep), solvation (fa_sol, lk_ball_wtd), electrostatics (fa_elec), hydrogen bonding energies (hbond_sc, hbond_sr_bb, hbond_lr_bb), and inter-chain interaction metrics (interchain_vdw, interchain_pair).

For FlexPepDock outputs, 14 structure-based scoring terms were used, including total and reweighted scores (score, reweighted_sc), interface score (I_sc), buried surface area (I_bsa), hydrogen bonds (I_hb, hbond_sc), packing and unsatisfied polar contacts (I_pack, I_unsat), van der Waals terms (fa_atr, fa_rep), solvation (fa_sol), and structural deviation (rmsALL).

All features were standardized prior to PCA. Clustering of the top models was then performed using K-means (k = 4), enabling classification of peptides into distinct binding profiles.

### Data accessibility

The mass spectrometry proteomics data have been deposited to the ProteomeXchange Consortium via the PRIDE partner repository with the dataset identifier. All relevant RNA seq data are available from the Gene Expression Omnibus (GEO) under accession number.

Additional processed data and analysis scripts supporting the findings of this study are available upon reasonable request.

## Discussion

In this study, we identified and functionally validated a tumor-derived non-canonical peptide, APAPAPAPL, that exhibits specific binding to HLA-E and selectively activates aNK cells.

Through integrative mass spectrometry and transcriptomic analysis, combined with structural modeling via FlexPepDock and RosettaDock, we showed that APAPAPAPL forms a highly stable complex with HLA-E and CD94-NKG2C. Our findings provide compelling evidence that certain tumor-expressed, non-canonical peptides can serve as potent immunogenic cues and tumor vaccine for memory-like NK cell responses.

Tumor vaccines have emerged as a promising tool in cancer prevention and treatment, with personalized strategies increasingly applied in clinical settings. Broadly, tumor vaccines can be categorized into prophylactic and therapeutic types ^[47, 48]^. Prophylactic cancer vaccines are primarily designed to prevent virus-associated malignancies. Notable successes include the Human Papillomavirus (HPV) vaccine, which has dramatically reduced the incidence of HPV-induced cervical cancer ^[49]^, and the Hepatitis B virus (HBV) vaccine, which significantly lowers the risk of hepatocellular carcinoma (HCC) ^[50]^. Epstein-Barr Virus (EBV), a known risk factor for several malignancies—such as nasopharyngeal carcinoma (NPC), Burkitt lymphoma, Hodgkin lymphoma, NK/T-cell lymphoma, and certain gastric cancers ^[51, 52]^, has prompted growing efforts to develop effective EBV vaccines^[53, 54]^ . A recent study demonstrated that a novel EBV vaccine formulation incorporating a lymph node-targeting Amphiphile-CpG adjuvant, gp350 glycoprotein, and a polyepitope protein induces potent protective immune responses ^[55]^ Additionally, EBV mRNA vaccines have shown potential in enhancing NK cell-mediated clearance of NPC^[56]^. In contrast, therapeutic cancer vaccines primarily target endogenous tumor antigens and can be broadly categorized into two types: off-the-shelf vaccines and personalized vaccines. Given the substantial inter-patient heterogeneity in tumor antigen profiles, current development efforts are increasingly shifting toward personalized vaccine strategies.

The extensive diversity of immune receptors (TCRs and BCRs) and the polymorphism of MHC molecules provide the immunological foundation for tumor vaccine development, allowing either shared or personalized tumor antigens to be recognized and targeted across different patients, though effective antigen presentation remains a determinant.

Unlike the tumor antigens presented by polymorphic MHC molecules as a basis of T cells immunity, HLA-E presents a restricted repertoire due to its limited polymorphism. Most of the peptides HLA-E recognizes and presents are the leading peptides of MHC-I molecules^[57]^, and interacts with inhibitory NKG2A and activating NKG2C^[39, 58]^. HLA-E presents MHC-I-derived signal peptides to the inhibitory receptor NKG2A which has the high affinity in such a physiological state, maintaining NK cell tolerance under normal conditions. When MHC-I expression is reduced, inhibitory signals decline, leading to NK cell activation. Under infectious conditions—particularly during CMV infection—certain peptides presented by HLA-E exhibit higher affinity for the activating receptor NKG2C than for the inhibitory receptor NKG2A ^[30]^, thereby delivering activating signals to NK cells. In addition, CMV-derived peptides can promote the expansion of NKG2C^+^ NK cells in patients, contributing to the dominance of the NKG2C-mediated response ^[59, 60]^. Therefore, it is reasonable to hypothesize that certain tumor-derived antigens may mimic CMV peptides by binding to HLA-E and preferentially engaging the activating receptor NKG2C over the inhibitory NKG2A. This property positions them as promising candidates for the development of NK cell-targeted tumor vaccines, which forms the core of this study.

However, the limited polymorphism of HLA-E and NKG2C presents a challenge that may constrain the breadth of their clinical applicability as the basis of tumor vaccine. Nevertheless, the intrinsic conformational flexibility of these molecules offers an opportunity. With advances in protein *de novo* design, it is now feasible to engineer specific stabilizers or binders that enhance the stability of otherwise weak peptide-HLA-E complexes and promote their preferential recognition by NKG2C. This strategy may significantly expand the range of tumor antigens effectively targeted by NK cells. On the other hand, identifying peptides that can be simultaneously recognized by both NK cells and T cells, through analyzing shared sequence or structural characteristics, offers another promising strategy. Such bi-recognizable peptides may enable coordinated activation of innate and adaptive immunity, thereby enhancing the overall antitumor immune response through synergistic stimulation of NK and T cell pathways.

## Acknowledgement

Sequencing was performed by the SNP&SEQ Technology Platform in Uppsala. The facility is part of the National Genomics Infrastructure (NGI) Sweden and Science for Life Laboratory. The SNP&SEQ Platform is also supported by the Swedish Research Council and the Knut and Alice Wallenberg Foundation.

## Author contributions

DS and YS proposed the hypothesis that adaptive NK cells recognize tumor-derived peptides via NKG2C and jointly designed the *in vitro* and *in vivo* assays. YS performed peptide experiments, single-cell RNA sequencing, and protein docking, while YK, RMB, JL and KG developed and analyzed the HLA-E immunoprecipitation and RNA sequencing strategies, with support from WS, SL, OG, and SS. YS and DS drafted the manuscript with input from all authors, and DS managed journal correspondence.

---

**Supplementary Figure 1.**
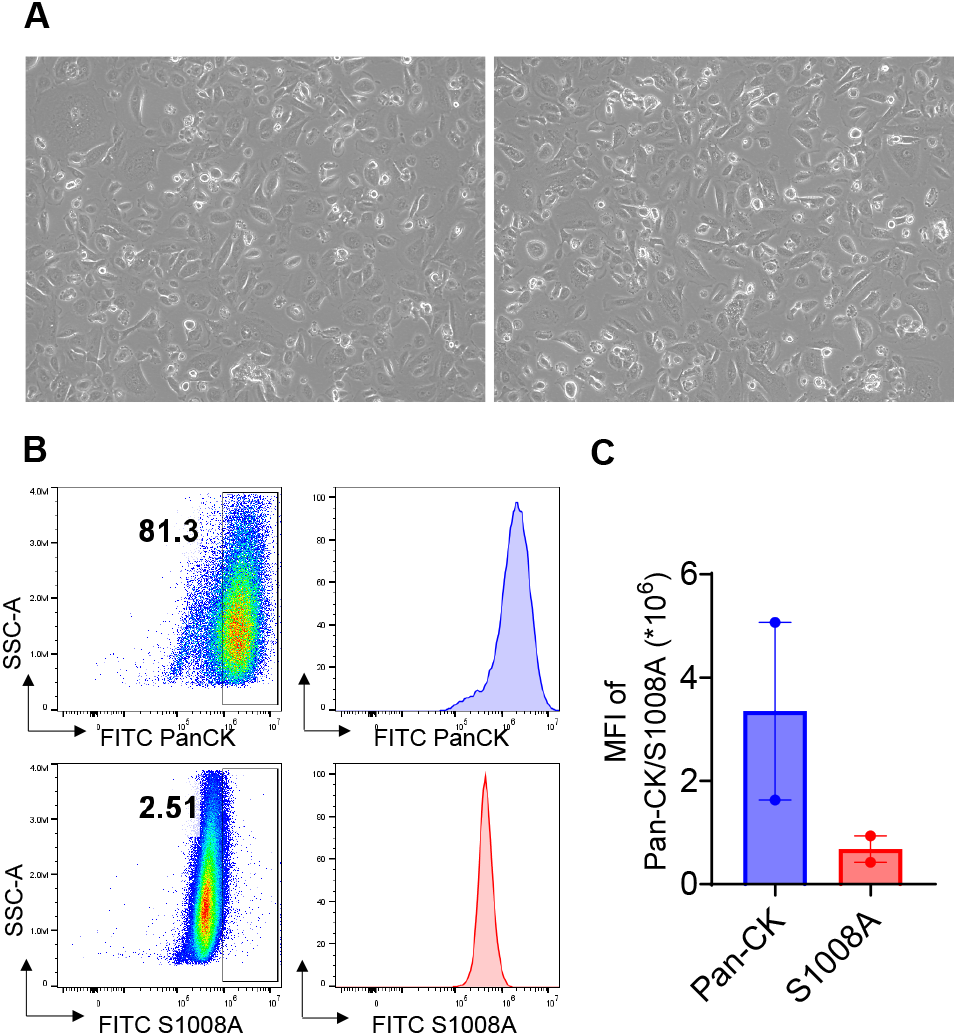
Validation of IPLA primary tumors. **(A)** Microscopic morphology of IPLA primary tumors. **(B)** Representative flow cytometry plots showing Pan-CK and S100A8 staining of IPLA primary tumors. **(C)** Summary data presented as mean ± SEM (n = 2).

**Supplementary Figure 2.**
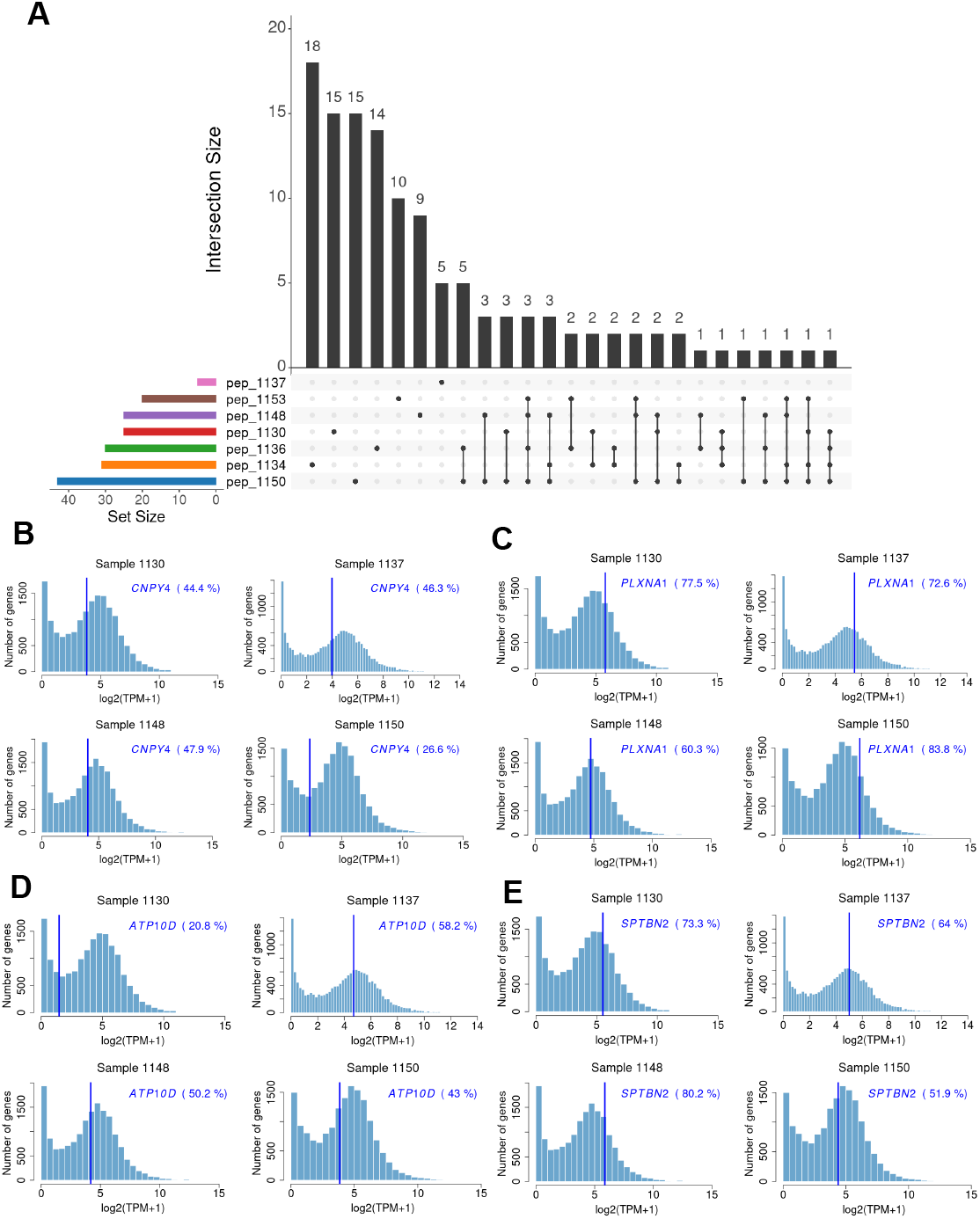
Shared HLA-E–bound peptides and transcriptomic validation of selected peptide-encoding genes. **(A)** UpSet plot illustrating shared and unique peptides among representative HLA-E–bound peptide repertoires. Horizontal bars show the total number of peptides identified per sample (set size), while vertical bars indicate the size of intersections across different sample combinations. **(B-E)** Histograms display the log_2_-transformed TPM expression distribution of all genes in each patient sample (1130, 1137, 1148, and 1150). Vertical blue lines indicate the position of the selected gene (italicized labels), and the corresponding percentile rank of the gene relative to all expressed genes in that sample is shown in parentheses. Panels illustrate representative examples for *CNPY4* **(B)**, *PLXNA1* **(C)**, *ATP10D* **(D)**, and *SPTBN2* **(E)**.

**Supplementary Figure 3.**
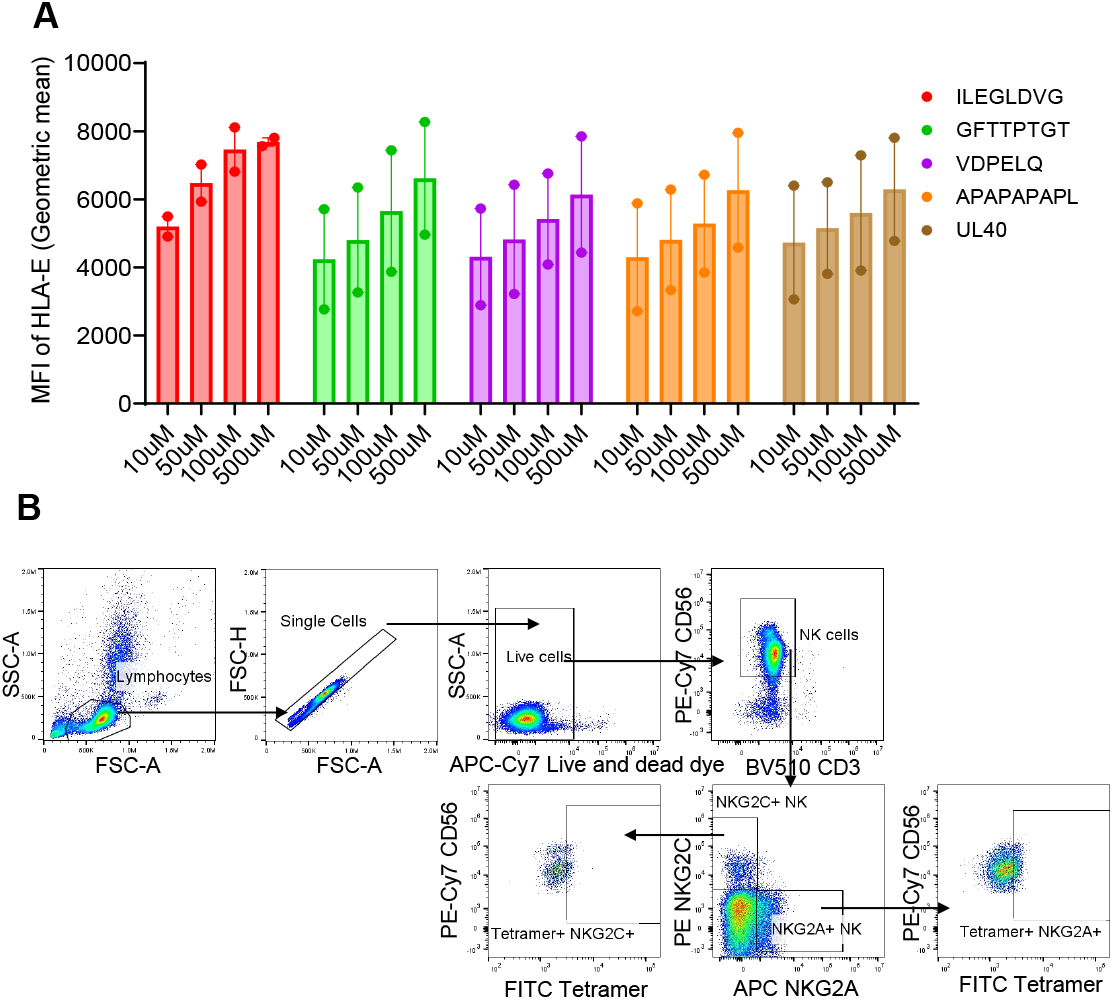
Titration for peptide concentration for HLA-E stabilization assay and gating strategies for Figure 2. **(A)** Frequencies of HLA-E on K562-HLA-E*0101 cells under different concentrations of peptides (n=2). **(A)** Representative flow cytometry gating strategy for tetramer staining: lymphocytes were gated on FSC/SSC, followed by single-cell and live-cell selection, CD3^−^CD56^+^ NK cell identification, and further subdivision into NKG2C^+^ or NKG2A^+^ NK subsets. Tetramer binding within NKG2C^+^ and NKG2A^+^ NK subsets is shown.

**Supplementary Figure 4.**
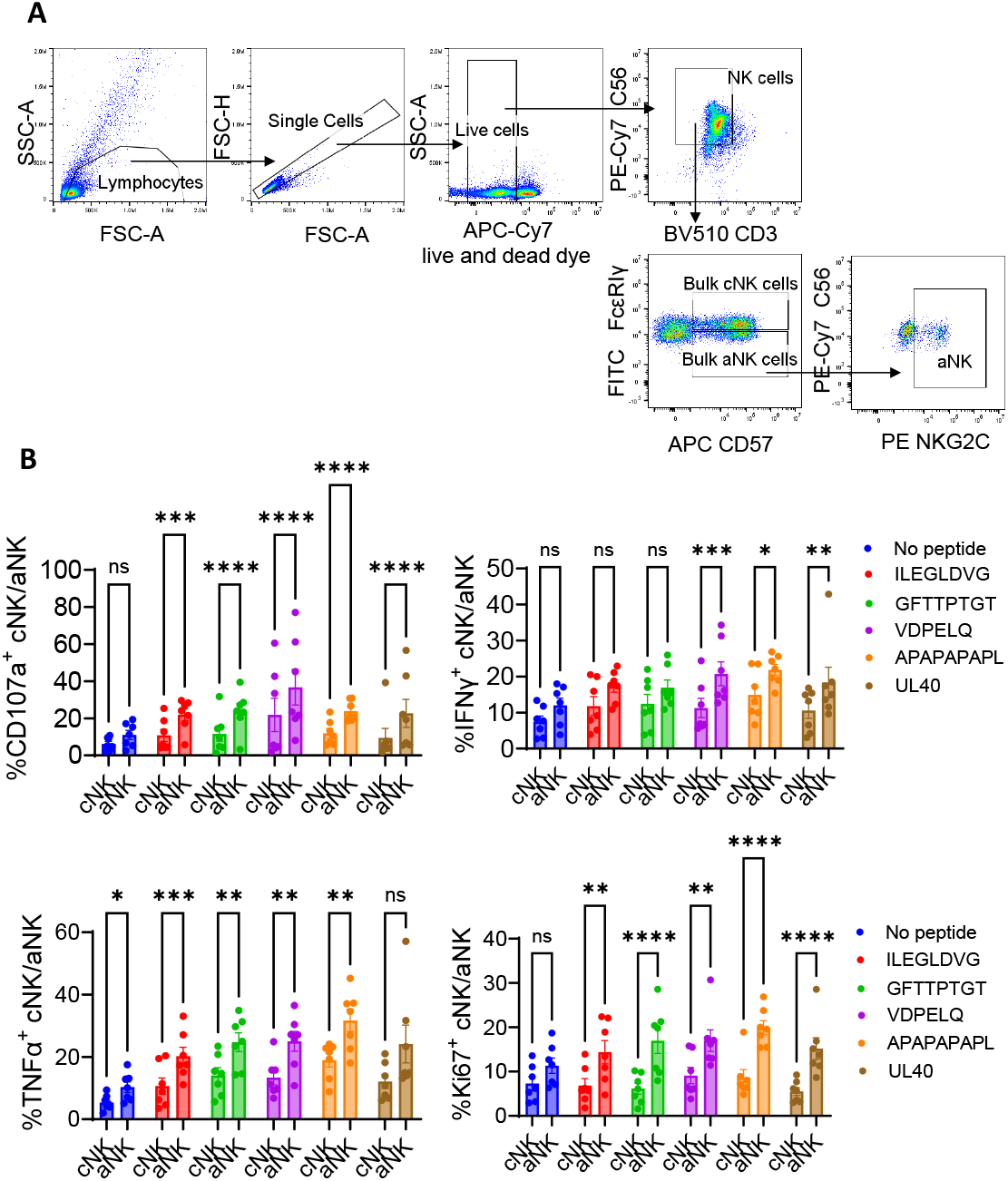
aNK cells elicit stronger first responses compared to cNK cells. **(A)** Flow cytometry gating strategy used to define NK-cell subsets. Lymphocytes were gated by FSC/SSC, followed by single-cell and live-cell selection. NK cells were identified as CD3^−^CD56^+^, then further subdivided into bulk cNK and aNK populations. (B) Summary of cNK and aNK first responses after 6 h co-culture with autologous DCs pulsed with the indicated peptides in terms of degranulation, cytokine production and proliferation (CD107a, IFN-γ, TNF-α, and Ki67) across donors (n = 7). Data are mean ± SEM; statistics by a two-way ANOVA (*p < 0.05, **p < 0.01, ***p < 0.001, ****p < 0.0001).

**Supplementary Figure 5.**
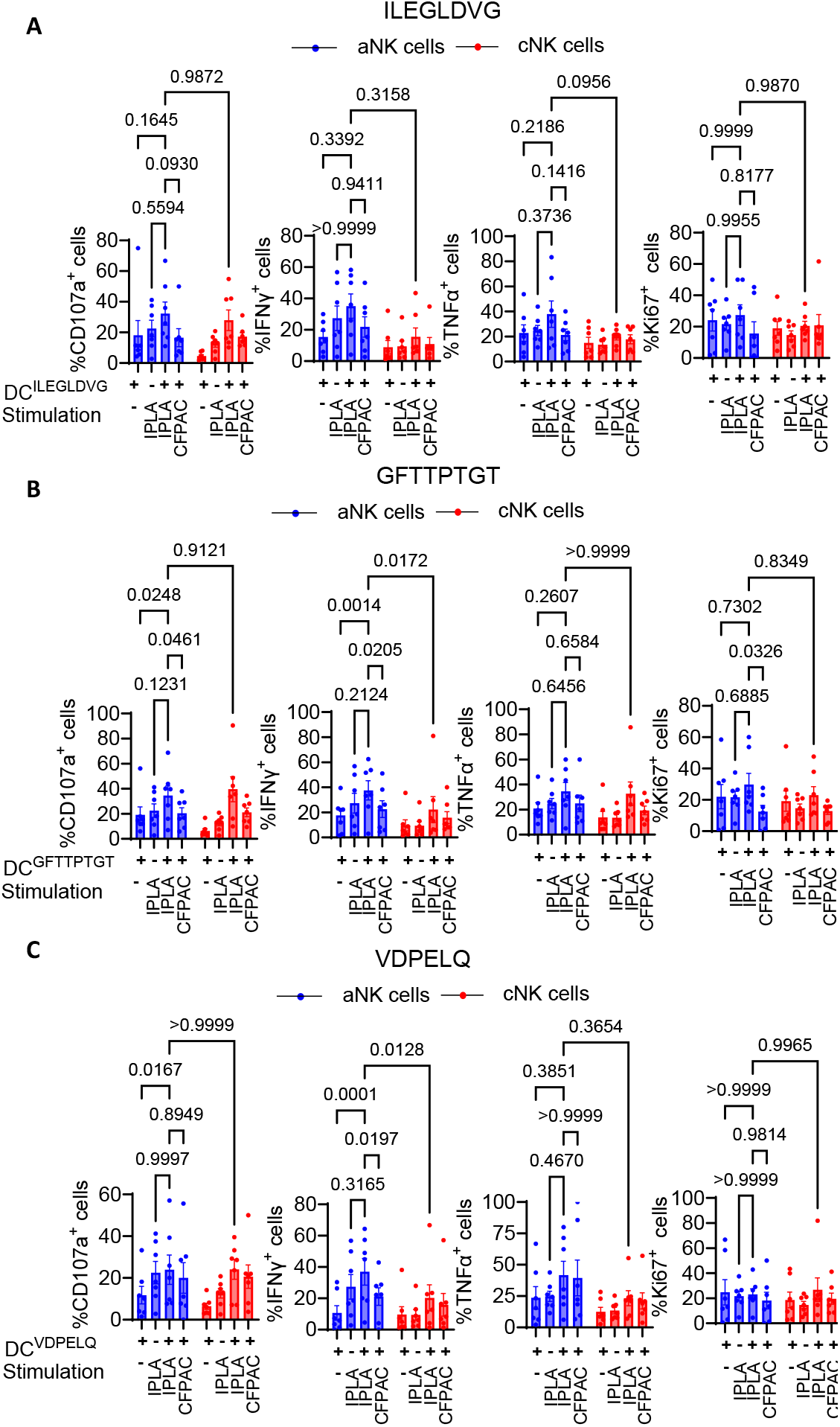
Three peptides (ILEGLDVG, GFTTPTGT, VDPELQ) does not elict the a NK recall responses very significantly. **(A-C)** Summary of of aNK and cNK cells upon IPLA tumor restimulation after 3-week co-culture with autologous DCs pulsed with ILEGLDVG **(A)**, GFTTPTGT **(B)** and VDPELQ **(C)**, showing degranulation, cytokine production, and proliferation (CD107a, IFN-γ, TNF-α, Ki67). aNK responses across donors (n = 7). Data are mean ± SEM; statistics by a two-way ANOVA with p-value indicated

**Supplementary Figure 6.**
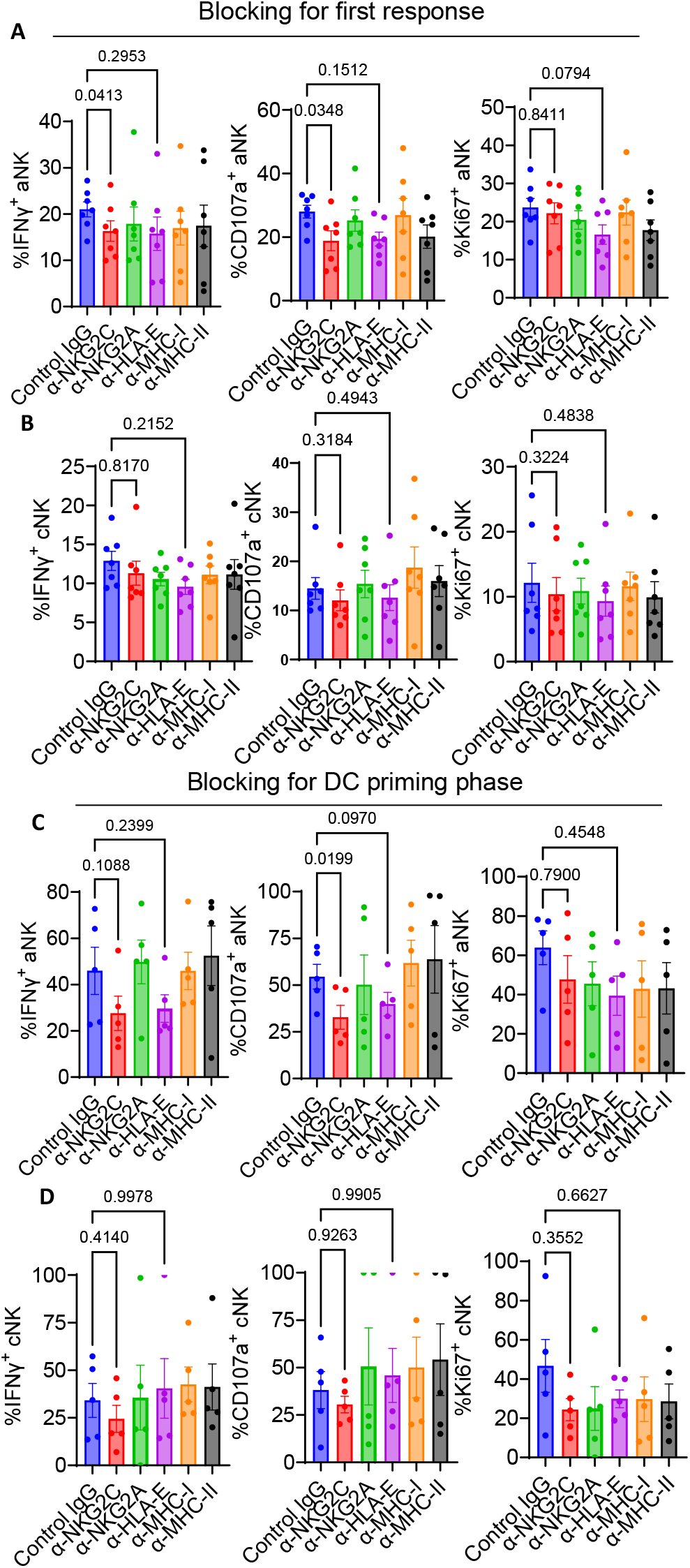
NKG2C and HLA-E are critical for first and recall responses. **(A-B)** Statistical summaries of IFN-γ^+^, CD107a^+^, and Ki67^+^ aNK / cNK cells with blocking antibodies after 6h coculture with DCs compared to isotype IgG control (n = 7). **(C-D)** Statistical summaries of IFN-γ^+^, CD107a^+^, and Ki67^+^ aNK / cNK cells with priming-phase blocking antibodies after 3-week coculture with DCs compared to isotype IgG control (n = 5). Data are mean ± SEM, one-way ANOVA conducted with p-value indicated

**Supplementary Figure 7.**
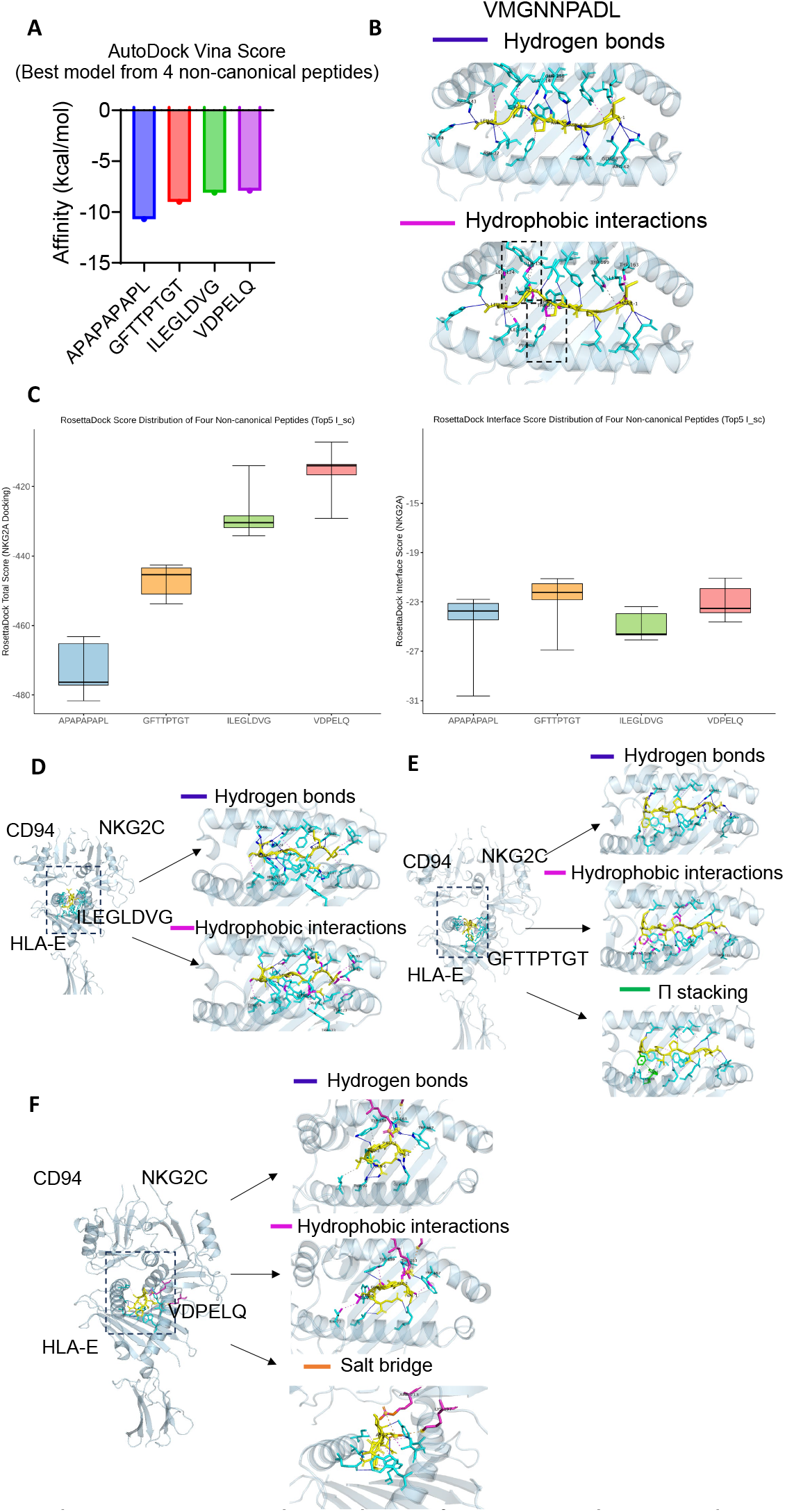
*In silico* simulations of APAPAPAPL-analogous peptide and other non-canonical peptides interacting with HLA-E and the CD94–NKG2C receptor. **(A)** AutoDock Vina affinity scores for four non-canonical peptides. **(B)** Predicted interaction sites between VMGNNPADL and HLA-E. **(C)** RosettaDock analysis of four non-canonical peptide–HLA-E complexes with CD94–NKG2A, with box plots (boxes = IQR, line = median, whiskers = range) showing total score and interface ΔG (I_sc) from the top five models. **(D–F)** Representative interaction maps of ILEGLDVG, GFTTPTGT, and VDPELQ bound to HLA-E in complex with CD94–NKG2C.

**Supplementary Figure 8.**
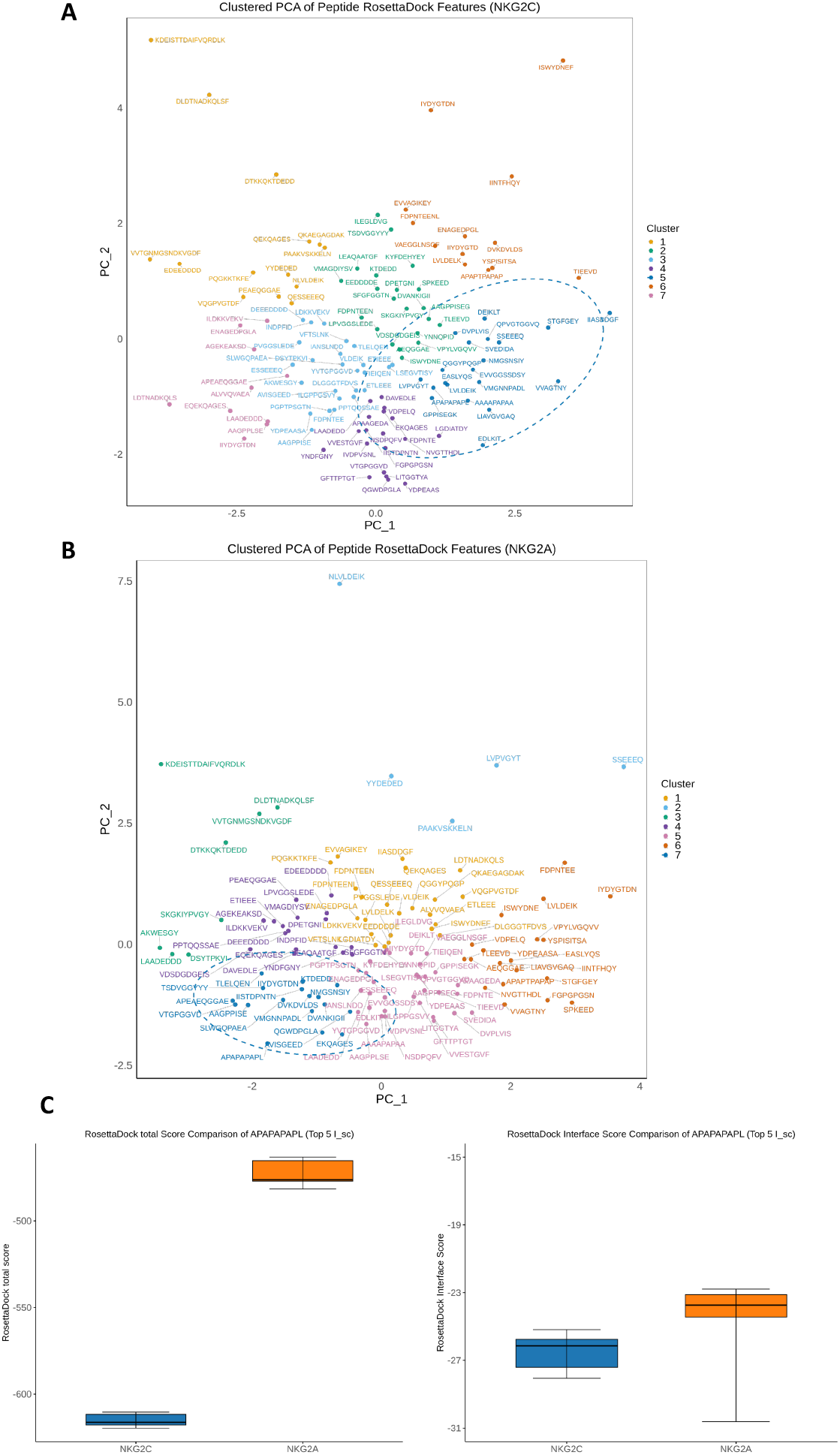
Comparative binding stability of CD94–NKG2C versus CD94–NKG2A to APAPAPAPL. **(A)** PCA mapping of all best models of peptides-HLA-E complexes interacting with CD94-NKG2C, showing their clustering based on multiple interaction simulation parameters. **(B)** PCA mapping of all best models of peptides-HLA-E complexes interacting with CD94-NKG2A, showing their clustering based on multiple interaction simulation parameters. **(C)** RosettaDock analysis comparing APAPAPAPL–HLA-E complexes with CD94–NKG2C or CD94–NKG2A. Box plots (boxes = interquartile range, line = median, whiskers = range) summarize the total score and interface ΔG (I_sc) from the top five models.

